# Survivin is a mechanosensitive cell cycle regulator in vascular smooth muscle cells

**DOI:** 10.1101/2022.11.09.515885

**Authors:** John C. Biber, Andra Sullivan, Joseph A. Brazzo, Amanda Krajnik, Yuna Heo, Kerry E. Poppenberg, Vincent M. Tutino, Su-Jin Heo, John Kolega, Kwonmoo Lee, Yongho Bae

## Abstract

Stiffened arteries are a pathology of atherosclerosis, hypertension, and coronary artery disease and a key risk factor for cardiovascular disease events. The increased stiffness of arteries triggers the hypermigration and hyperproliferation of vascular smooth muscle cells (VSMCs), leading to neointimal hyperplasia and accelerated neointima formation, but the mechanism of this trigger is not known. Our analyses of whole-transcriptome microarray data sets from mouse VSMCs cultured on stiff hydrogels simulating arterial pathology and from injured mouse femoral arteries revealed 80 genes that were differentially regulated (74 upregulated and 6 downregulated) relative to expression in control VSMCs cultured on soft hydrogels and in uninjured femoral arteries. A functional enrichment analysis revealed that these stiffness-sensitive genes are linked to cell cycle progression and proliferation. Furthermore, we found that survivin, a member of the inhibitor of apoptosis protein family, mediates stiffness-sensitive cell cycling and proliferation *in vivo* and *in vitro* as determined by gene network and pathway analyses, RT-qPCR, and immunoblotting. The stiffness signal is mechanotransduced via FAK and Rac signaling to regulate survivin expression, establishing a regulatory pathway for how the stiffness of the cellular microenvironment affects VSMC behaviors. Our findings indicate that survivin is necessary for VSMC cycling and proliferation and regulates stiffness-responsive phenotypes.

## INTRODUCTION

Arterial stiffening contributes to the development and progression of a variety of cardiovascular diseases [1–9]. The change in arterial stiffness affects the biophysical input to resident vascular smooth muscle cells (VSMCs), which constitute the biomechanically active and dynamic cell layer in the media of arteries. An increase in arterial stiffness triggers VSMCs to transition from a contractile (or differentiated) state to a synthetic (or dedifferentiated) state, in which they aberrantly migrate, proliferate, and produce extracellular matrix (ECM). This results in pathological neointima formation and further arterial stiffening [8–11]. The stiffness of the ECM surrounding VSMCs thus regulates cardiovascular biology with implications for aging [12] and Hutchinson–Gilford progeria syndrome [13] as well as atherosclerosis [5, 14], hypertension [15, 16], coronary artery diseases [17], and fibrosis [18].

To explore how ECM stiffness affects VSMC function, we and others have used fibronectin- or collagen-coated deformable polyacrylamide hydrogels to create culture matrices mimicking normal (healthy) and pathological (diseased) stiffnesses. Studies with this system showed that ECM stiffness affects cell cycling and proliferation [4, 7, 19], migration [20–22], intracellular stiffness and traction force [7, 19, 23, 24], cell–cell adhesion [19], and ECM synthesis [5, 12]. However, the signaling pathways regulating the response of VSMCs to ECM stiffness have yet to be fully defined.

There is evidence that VSMCs response to vascular injury and stiffening involve survivin [25–27]. Survivin (also known as BIRC5) belongs to the inhibitor of an apoptosis protein (IAP) family and was initially described as an antiapoptotic protein in cancer but later shown to regulate cancer cell migration and proliferation [28, 29]. Survivin is expressed at a low level in healthy adult tissue but is rapidly upregulated in response to vascular injury, atherosclerosis, and hypertension (conditions in which arteries stiffen [4, 5, 7]) in animal models [25, 26, 30]. Moreover, survivin is highly upregulated in proliferating VSMCs in the neointima and media in human atherosclerotic plaques and stenotic vein grafts [26]. Interestingly, overexpression of survivin (via a dominant-negative form of survivin) reduces neointima formation in rabbits [25], suggesting that survivin is a regulator of these effects.

Survivin induction after vascular injury correlates with the expression of cell proliferation genes downstream of focal adhesion kinase (FAK), which activates Rac to promote stiffness-sensitive cell cycle progression of VSMCs [7]. Furthermore, inhibition of FAK [31] or Rac [32] reduces survivin levels in other cell types. Because FAK and Rac are activated at sites of vascular injury and in VSMCs cultured on stiff hydrogels [4, 7, 19], we hypothesized that they contribute to the mechanotransduction of ECM stiffness to induce VSMC responses mediated by survivin. The objective of the present study was to identify the regulatory pathway through which the stiffness of the cellular environment affects VSMC behavior. Our findings suggest that ECM stiffness coordinates with survivin and focal adhesion biology in an integrated mechanobiochemical system to control the cell cycle progression and proliferation of VSMCs.

## RESULTS

### Whole-transcriptome analyses identify the stiffness-sensitive transcriptome of VSMCs

We examined the impact of ECM stiffness on the transcriptome of VSMCs *in vivo* and *in vitro.* For the *in vivo* study, we analyzed previously acquired whole-transcriptome microarray data [7] to identify genes that are differentially expressed after fine-wire injury to the femoral artery of mice (an *in vivo* model of VSMC proliferation and arterial stiffening). For the *in vitro* study, a whole-transcriptome microarray analysis was performed using mRNA samples from mouse VSMCs (mVSMCs) cultured for 24 h on fibronectin-coated soft (2–4 kPa) or stiff (16–30 kPa) polyacrylamide hydrogels, which reflect the elastic moduli of healthy or injured/diseased mouse artery [4, 7, 19]), respectively. Our *in vivo* transcriptome data analysis identified 25,253 genes (11,336 upregulated and 13,917 downregulated genes in injured femoral arteries compared to expression in uninjured arteries) (**Fig. 1A**), and the *in vitro* data analysis identified 21,999 genes (9,627 upregulated and 12,372 downregulated genes in mVSMCs cultured on stiff hydrogels compared to expression in cells on soft hydrogels) (**Fig. 1B**).

**Figure 1.**
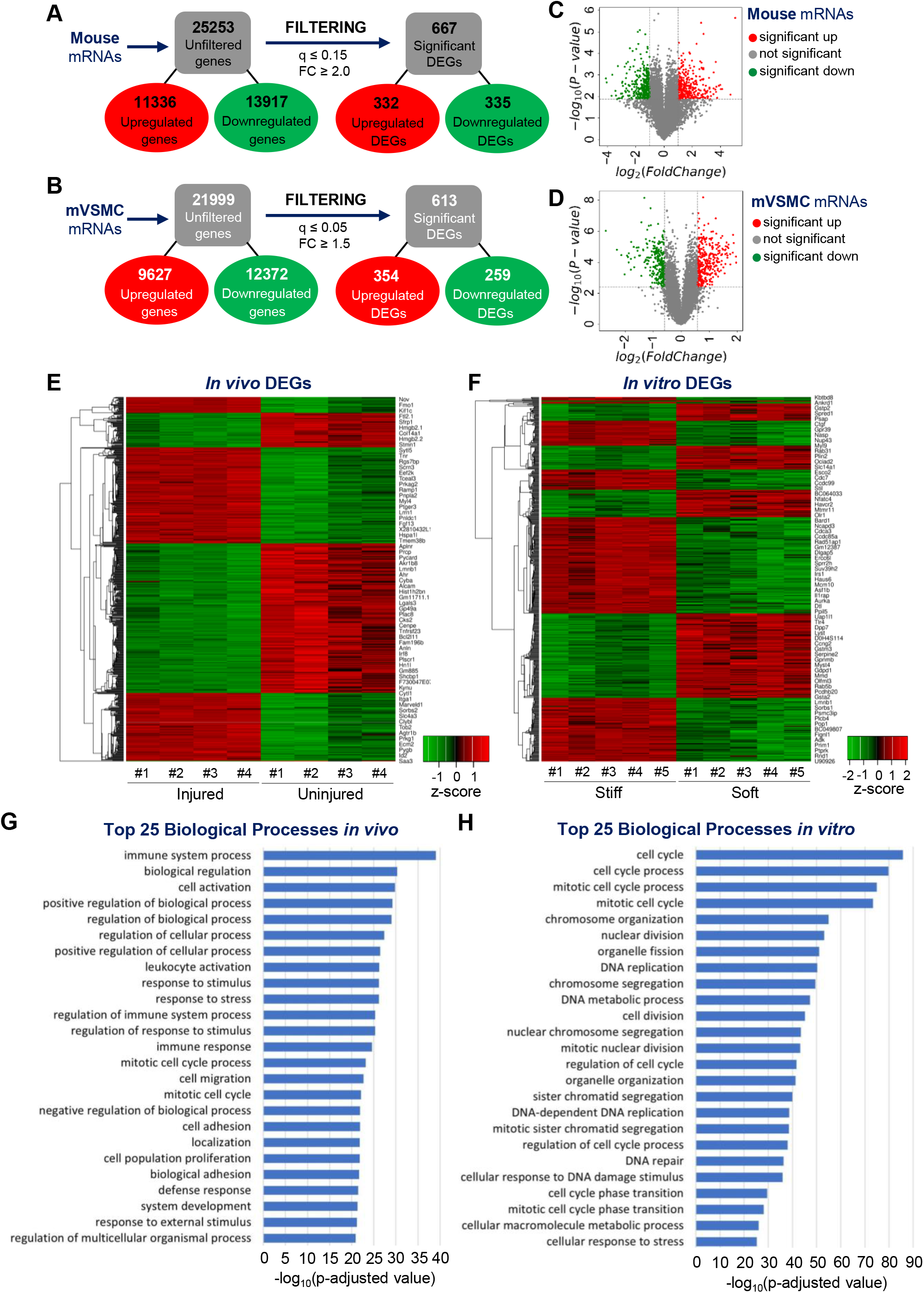
Whole-transcriptome analyses show the stiffness-sensitive transcriptome of vascular smooth muscle cells (VSMCs). Reductions in data magnitude by applying significance thresholds to the raw *in vivo* data (fold change [FC] ≥ 2.0, *q* value ≤ 0.15) (**A**) and the raw *in vitro* data (FC ≥ 1.5, *q* value ≤ 0.05) (**B**). Volcano plots display the distributions of all detected transcripts, represented as single dots that are not statistically different (gray), significantly upregulated (red), or significantly downregulated [33] from *in vivo* (**C**) and *in vitro* (**D**) data sequencing results. The *y* axis represents each gene’s −log_10_(*p* value), whereas the *x* axis represents their log_2_(fold change). Heat maps display the Z-scores of the 667 differentially expressed genes (DEGs) *in vivo* (**E**) and 613 DEGs *in vitro* (**F**). Histograms present the top 25 biological processes enriched for 332 upregulated DEGs in the *in vivo* data set (**G**) and 354 upregulated DEGs in the *in vitro* data set (**H**).

These data were then filtered by fold change and *q* value (see Materials and Methods for details), resulting in a list of 667 statistically significant differentially expressed genes (DEGs) for the *in vivo* study (332 upregulated and 335 downregulated; **Fig. 1A** and **Table S1**) and 613 significant DEGs for the *in vitro* study (354 increased and 259 decreased; **Fig. 1B** and **Table S2**). The distributions of these DEGs against the total number of identified genes were plotted as the −log_10_(*p* value) versus log_2_(fold change) values of each detected gene for the *in vivo* (**Fig. 1C**) and *in vitro* (**Fig. 1D**) data sets, with red color denoting significantly upregulated DEGs, green denoting significantly downregulated DEGs, and gray denoting genes with no significant change. Additionally, the DEGs obtained from *in vivo* (**Table S1**) and *in vitro* (**Table S2**) data sets are depicted in heat maps, showing significant clustering (uninjured vs. injured femoral arteries [**Fig. 1E**] and mVSMCs on soft vs. stiff hydrogels [**Fig. 1F**]).

We conducted a gene ontology (GO)-based functional enrichment analysis to identify biological processes associated with the upregulated DEGs. *In vivo* DEGs were highly enriched in cell activation, regulation of biological/cellular processes, response to stress or stimulus, mitotic cell cycle process, migration, and proliferation (**Fig. 1G**), all of which are important in neointima formation associated with arterial stiffening. DEGs upregulated *in vitro* were highly enriched primarily in the regulation of cell cycle processes (**Fig. 1H**), which are critical for cell cycle progression and proliferation.

### Network analysis identifies survivin as a stiffness-sensitive mediator of cell cycle progression and proliferation

There were 80 DEGS common between the *in vitro* and *in vivo* datasets: 74 genes were commonly upregulated (**Fig. 2A**) and 6 genes were commonly downregulated (**Fig. 2B**). The expression levels of these common DEGs are presented as heat maps with hierarchical clustering, demonstrating groups of genes whose relative expression changes were most similar in the two data sets (**Fig. 2C, D**). We used Cytoscape’s String application to couple biological processes with a network analysis of the 74 commonly upregulated DEGs. The functional enrichment of the DEGs in this string network indicated that the genes were involved in cell cycle regulation and nuclear/cell division (**Fig. 2E**). Notably, the most highly connected DEG among them was *Birc5* (baculoviral IAP repeat-containing 5; encodes survivin). *Birc5* was significantly upregulated in both *in vitro* and *in vivo* data sets and linked to all of the biological processes highlighted in this network. We therefore hypothesized that survivin serves as a molecular linchpin that controls the stiffness-mediated cell cycle progression of VSMCs.

**Figure 2.**
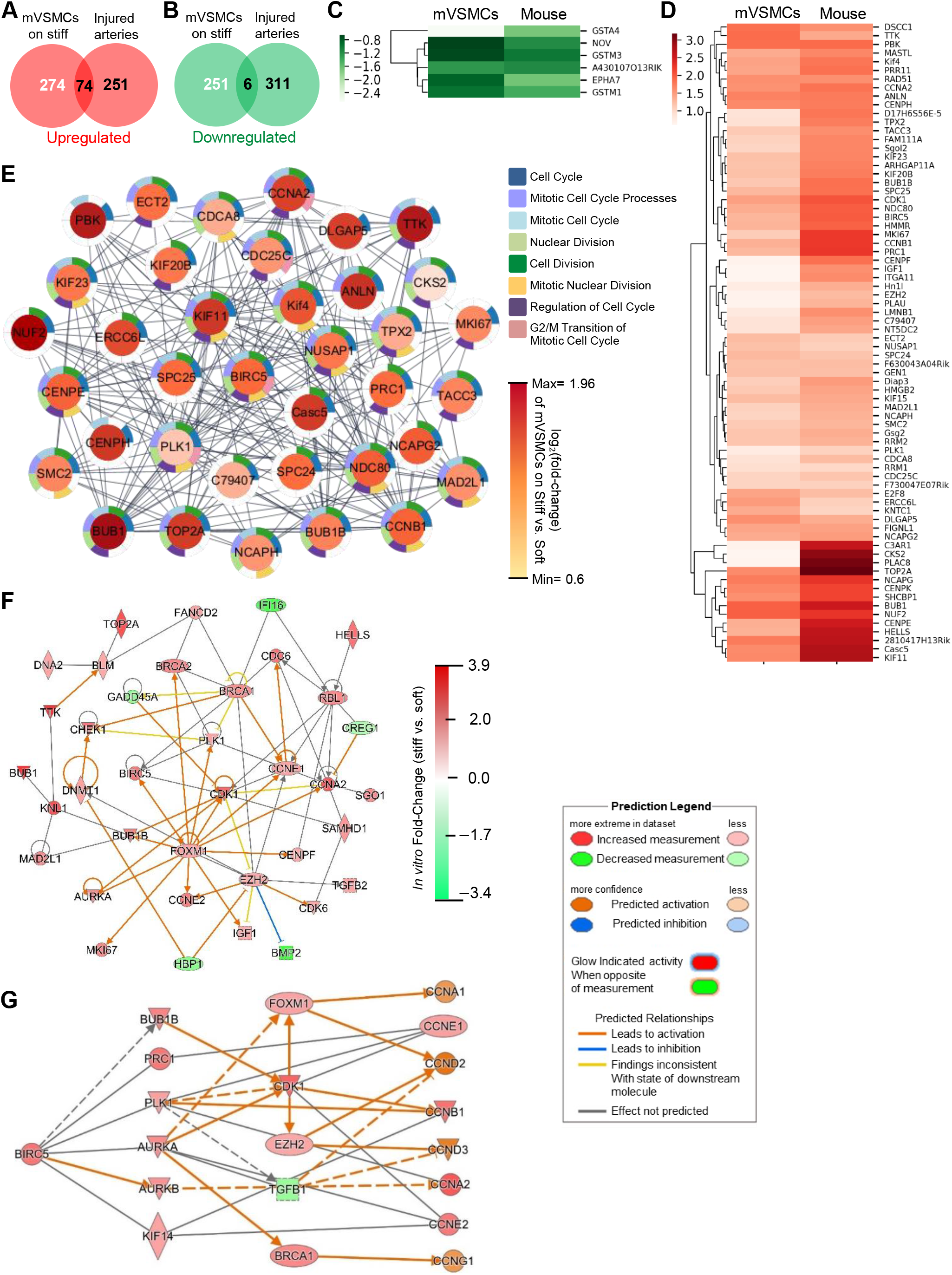
Network analysis identifies survivin *(Birc5)* as a potential stiffness-mediated regulator of cell cycle progression. Venn diagrams representing the number of commonly upregulated (**A**) or downregulated (**B**) differentially expressed genes (DEGs) present within the *in vivo* and *in vitro* data sets. Heat maps representing the log_2_(fold change) of the 6 DEGs that were commonly downregulated (**C**) and the 74 DEGs that were commonly upregulated (**D**) in both *in vivo* and *in vitro* data sets. Hierarchical clustering dendrograms are provided to the left of each heat map. (**E**) String network generated using the 74 commonly upregulated DEGs, with node color representing *in vitro* log_2_(fold change) values and border colors indicating DEG memberships to GO biological process categories. (**F**) Network diagram providing an overview of the molecular interactions related to Birc5 within the activated cell cycle progression function (Z-score = 2.124) in cardiovascular disease. (**G**) Predictive model for Birc5 regulation of cyclins. Network diagram presenting downstream targets of Birc5 (including cyclins) and potential intermediate molecules, with node color and intensity representing the observed expression levels or predicted activation states based on the *in vitro* data.

Ingenuity Pathway Analysis (IPA; Qiagen) was applied to the *in vitro* DEGs to identify genes that interact with *Birc5* and are involved with cell cycle progression (with a Z-score of 2.124; a Z-score of ≥2 is considered significant [7, 33] and associated with cardiovascular disease (**Fig. 2F**)), including *Ccna2, Ccne1,* and *Ccne2.* To further visualize the molecular relationships between *Birc5* and cell cycle-associated DEGs, we generated another network with fold change values overlayed from the *in vitro* data set (**Fig. 2G**) using the Path Explorer tool of the IPA program. The survivin-to-cyclin network with the *in vivo* data set (**Fig. S1**) was similar to that for the *in vitro* data.

### Survivin in VSMCs is stimulated by pathological ECM stiffness

An analysis of the *in vivo* and *in vitro* whole-transcriptome data sets for IAP family members other than *Birc5* (for survivin) identified *Naip1* (NLR family, apoptosis inhibitory protein 1 *[Birc1]), Birc2, Birc3, Xiap* (X-linked IAP *[Birc4]), Birc6* (baculoviral IAP repeat-containing 6), and *Birc7.*However, only *Birc5* mRNA expression was significantly increased in injured arteries (4-fold compared to that in the uninjured control; **Fig. 3A**) and in mVSMCs grown on stiff hydrogels (2.5-fold compared to that in cells grown on soft hydrogels; **Fig. 3B**). The increases in survivin levels were confirmed with real-time quantitative PCR (RT-qPCR) and immunoblotting from mVSMCs (**Fig. 3C, D**) as well as human VSMCs (hVSMCs) (**Fig. 3F, G**) cultured on soft and stiff hydrogels. Furthermore, protein levels of cyclins D1 and A (major targets of ECM stiffness-mediated signaling for cell cycle progression [4, 7, 19]) were sensitive to ECM stiffness in mVSMCs (**Fig. 3D, E**) and hVSMCs (**Fig. 3G, H**), respectively. Together with the results described above from the functional enrichment analysis (**Fig. 2E**), these data suggest that stiffness-sensitive expression of survivin is associated with cell cycle progression.

**Figure 3.**
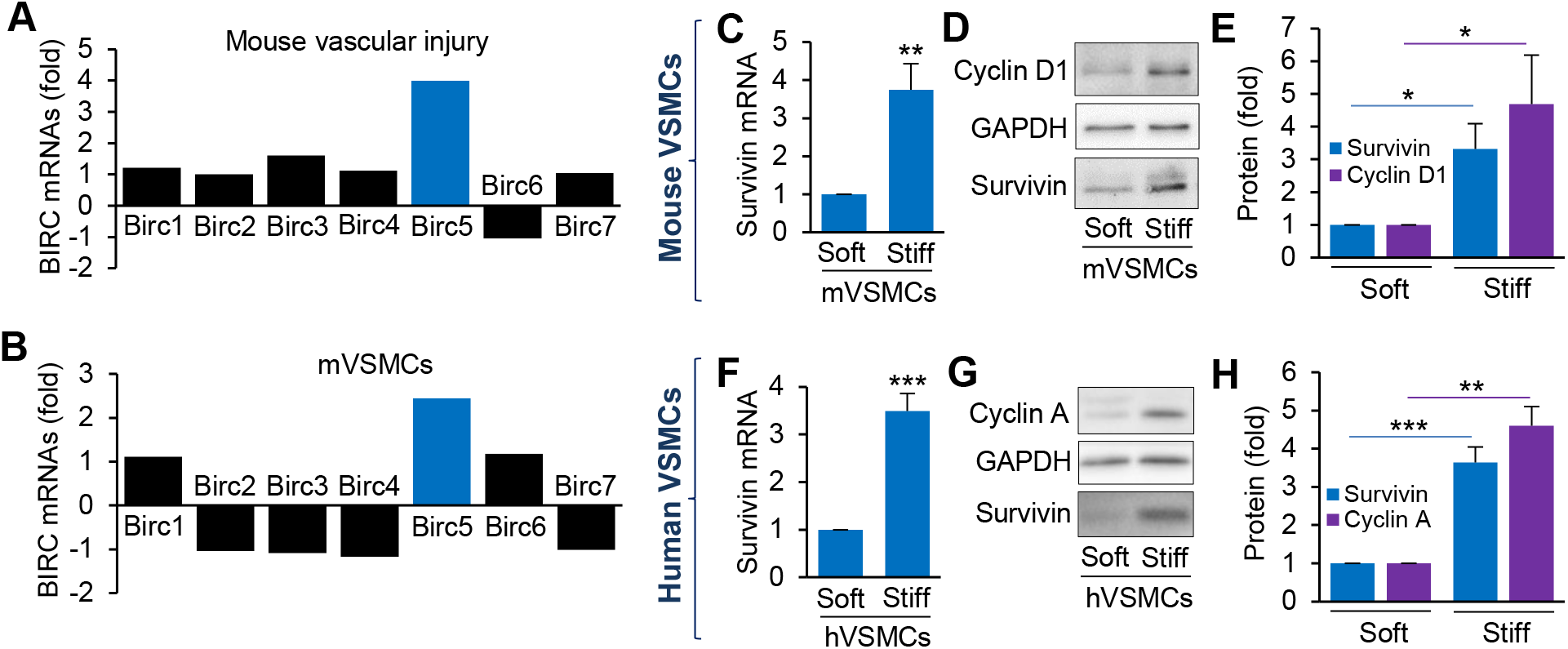
ECM stiffness modulates survivin expression in vascular smooth muscle cells (VSMCs). Differential mRNA expression of baculovirus inhibitor of apoptosis repeat-containing (BIRC) gene family after femoral vascular injury (**A**) and from mouse VSMCs (mVSMCs) seeded on fibronectin-coated soft or stiff polyacrylamide hydrogels (**B**). G_0_-synchronized mVSMCs (**C–E**) or human (h)VSMCs (**F–H**) were seeded on soft or stiff hydrogels for 24 h. Survivin gene (**C** and **F**) and protein (**D** and **G**) expression was analyzed by RT-qPCR and immunoblotting, respectively. Levels of survivin mRNA were normalized to those measured in mVSMCs (**C**) and hVSMCs (**F**) on soft hydrogels; *n = 5* (C) and *n = 8* (F) independent experiments. Average survivin intensity in mVSMCs (**D**) and hVSMCs (**G**) was quantified using ImageJ and normalized to that in VSMCs on soft hydrogels. Representative immunoblot images from four independent biological replicates; GAPDH served as a loading control. Survivin protein levels normalized to that in mVSMCs (**E**) and hVSMCs (**H**) on soft hydrogels. Data are means + SEMs. *\*p* < 0.05, \*\**p* < 0.01, \*\*\**p* < 0.001.

### Survivin is essential for stiffness-mediated cell cycle progression

We next investigated how survivin expression affects cell cycle progression and proliferation. We transfected hVSMCs with control or survivin siRNAs before culturing them on soft or stiff hydrogels. These analyses showed that the targeted siRNAs reduced survivin mRNA (**Fig. 4A**) and protein (**Fig. 4D, E**) expression by 60–70% in hVSMCs cultured on stiff hydrogels compared to that in cells with control siRNAs. Furthermore, knockdown of survivin decreased the stiffness-mediated induction of cyclin D1 (CCDN1; **Fig. 4B, F**) and cyclin A (CCNA; **Fig. 4C, G**). Accordingly, S-phase entry was reduced by survivin siRNAs in cells cultured on stiff hydrogels as determined by the incorporation of EdU (5-ethynyl-2’-deoxyuridine) (**Fig. 4H**). Similarly, blocking the induction of survivin with YM155 (a pharmacological agent that inhibits survivin expression) in cells cultured on stiff hydrogels (**Fig. S2A**) reduced expression of *Ccnd1* (**Fig. S2B**), *Ccna* (**Fig. S2C**), and *Ccnb* (cyclin B) (**Fig. S2D**). S-phase entry was triggered in hVSMCs plated on soft hydrogels with adenovirus-mediated overexpression of survivin (**Fig. 4I**) relative to that in hVSMCs infected with a control virus (for GFP expression). These data demonstrate that survivin in VSMCs mediates the effects of ECM stiffness on cell cycle progression.

**Figure 4.**
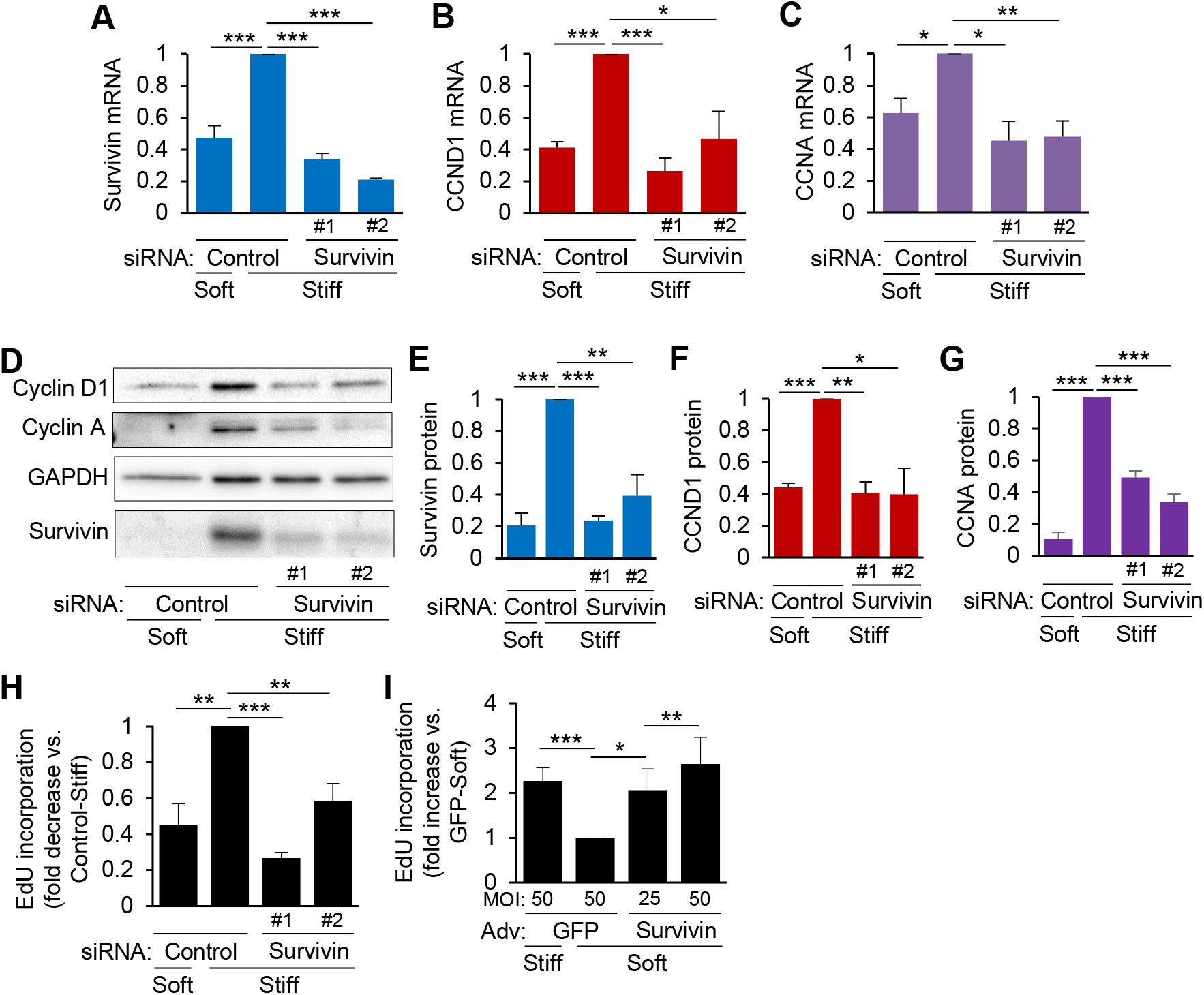
Survivin is required for stiffness-mediated cell cycle progression. (**A–G**) Human vascular smooth muscle cells (hVSMCs) were transfected with control siRNA or siRNAs to survivin [1 and #2], synchronized to G0 via serum starvation, and plated on soft or stiff hydrogels with 10% FBS for 24 h. Total cell lysates were analyzed by RT-qPCR (**A–C**) or immunoblotting (**D–G**) to determine mRNA and protein levels of survivin (**A** and **E**), cyclin D1 (CCND1; **B** and **F**), and cyclin A (CCNA; **C** and **G**). Expression levels were normalized to those in hVSMCs treated with control siRNA on stiff hydrogels. *n = 4* (A), *n = 3* (B), *n = 3* (C), *n = 5* (E), *n = 3* (F), and *n = 5* (G) independent experiments. S-phase entry with survivin knockdown and overexpression was assessed by EdU incorporation, which was normalized to hVSMCs treated with control siRNA on stiff hydrogels (**H**) or infected with GFP adenovirus (Adv) on soft hydrogels (**I**). *n = 5* (H) and *n = 5* (I) independent experiments. Data are means + SEMs. \**p* < 0.05, \*\**p* < 0.01, \*\*\**p* < 0.001.

### FAK-Rac signaling regulates stiffness-mediated survivin expression

Previous studies showed that the activity of FAK and Rac increases with vascular injury and stiffness-induced cell cycle progression and proliferation [4, 7, 19] and that survivin induction after vascular injury correlates with the expression of cell cycle genes downstream of FAK [7]. We therefore investigated whether FAK and Rac activity is responsible for stiffness-mediated survivin expression. Treatment of hVSMCs on stiff hydrogels with PF573228 (PF; FAK-specific inhibitor) or EHT1864 (EHT; Rac-specific inhibitor) markedly reduced the levels of survivin mRNA (**Fig. 5A**) and protein (**Fig. 5B, C**). These effects were confirmed by reduced survivin expression in hVSMCs infected with adenoviruses encoding FAK397 (a nonphosphorylatable form of FAK) or RacN17 (dominant-negative Rac) compared to that in cells expressing the LacZ control (**Fig. 5D**).

**Figure 5.**
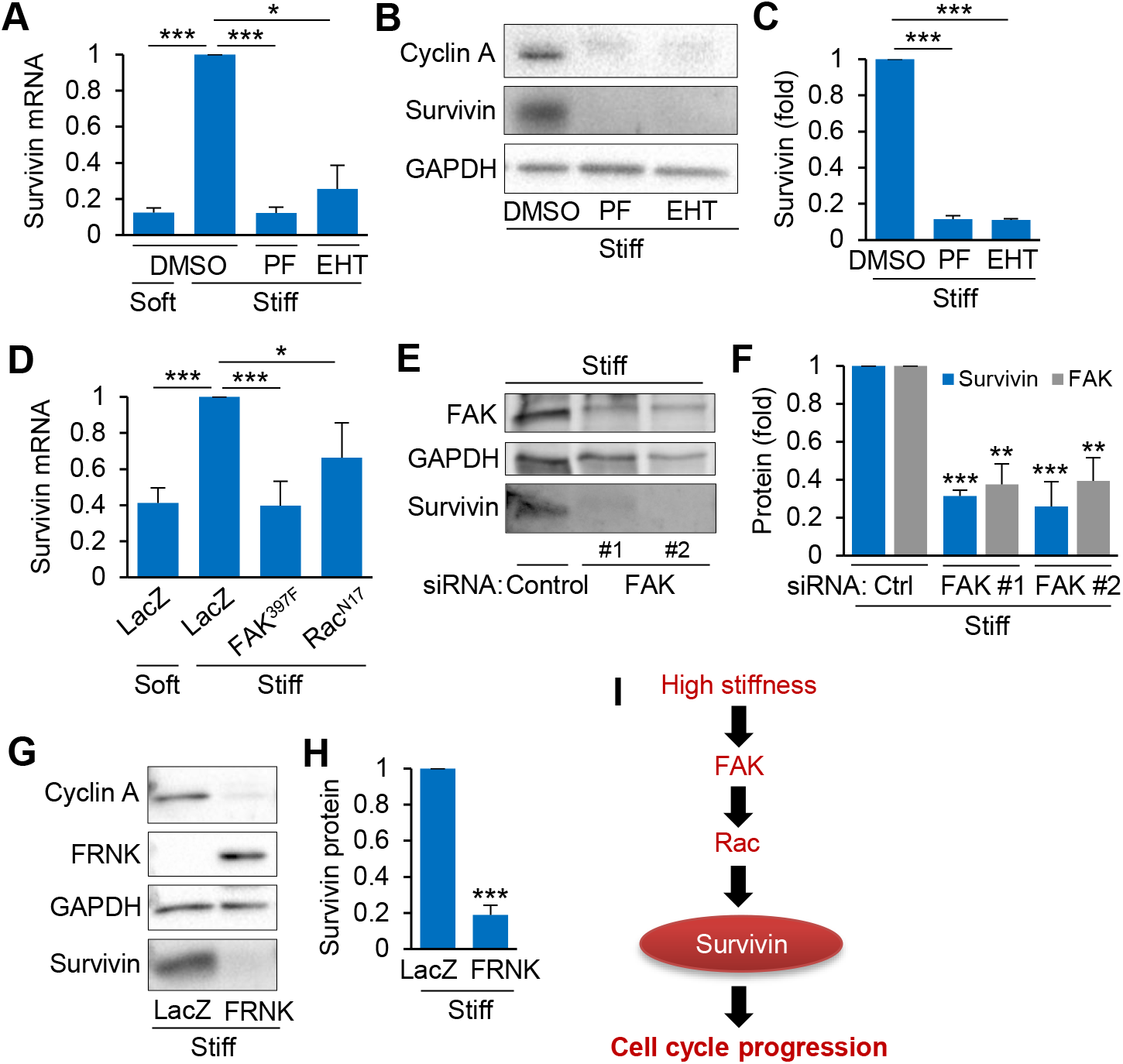
FAK-Rac signaling regulates stiffness-mediated survivin expression. G0-synchronized human vascular smooth muscle cells (hVSMCs) treated with FAK inhibitor PF573228 (PF) or Rac inhibitor EHT1864 (EHT) in dimethyl sulfoxide (DMSO) were seeded on fibronectin-coated soft or stiff hydrogels for 24 h. Survivin mRNA (**A**) and protein (**B** and **C**) expression was analyzed by RT-qPCR and immunoblotting, respectively; levels were normalized to those in hVSMCs treated with DMSO on stiff hydrogels; *n* = 3 (A) and *n* = 5 (B and C) independent experiments. (**D**) hVSMCs infected with adenoviruses encoding LacZ, FAK^397F^, or Rac^N17^ were seeded on soft or stiff hydrogels with 10% FBS for 24 h. Survivin mRNA expression was analyzed by RT-qPCR; levels were normalized to those in hVSMCs infected with LacZ on stiff hydrogels. *n* = 8 independent experiments. (**E** and **F**) G_0_-synchronized hVSMCs were transfected with control siRNA or siRNAs to FAK [1 and #2], serum-starved, and plated on stiff hydrogels with 10% FBS for 24 h. FAK and survivin levels in total cell lysates were analyzed by immunoblotting; levels were normalized to those in hVSMCs treated with control siRNA on stiff hydrogels. *n* = 3 independent experiments. (**G** and **H**) hVSMCs infected with adenoviruses encoding LacZ or FRNK were plated on stiff hydrogels for 24 h. Protein expression levels were analyzed by immunoblotting. *n* = 4 independent experiments. (**I**) Model of signal transduction for stiffness-mediated cell cycle progression. Data are means + SEMs. \**p* < 0.05, \*\**p* < 0.01, \*\*\**p* < 0.001.

Additionally, siRNA-mediated knockdown of FAK (**Fig. 5E, F**) and adenovirus-mediated overexpression of FAK-related non-kinase (FRNK) (**Fig. 5G, H**) similarly reduced survivin levels in hVSMCs cultured on stiff hydrogels compared to that in controls. Collectively, our results suggest that the mechanotransduction of ECM stiffness activates FAK-Rac signaling, which upregulates survivin and triggers cell cycle progression in VSMCs (**Fig. 5I**).

## DISCUSSION AND CONCLUSION

Stiffness-mediated remodeling of the vascular wall involves complex interactions between the local microenvironment and VSMCs. Moreover, accelerated cell cycling and proliferation of VSMCs appear to both cause and result from the arterial stiffening process. To identify the molecular mechanism by which stiffness affects VSMC proliferation, we applied integrative genome-wide analyses to databases from mouse models of vascular injury and cellular models of arterial stiffening and cell proliferation. The evidence suggests that survivin is the molecular linchpin by which arterial or ECM stiffness mediates pathological VSMC behaviors.

Survivin is known as a critical regulator of mitosis and cytokinesis during cancer cell division. Overexpression of survivin in the nuclei of cancer cells promotes the G1-S cell cycle transition by releasing p21 (known as a cyclin-dependent kinase [Cdk] inhibitor 1] from Cdk4 to activate the cyclin D1/Cdk4 complex, leading to the phosphorylation of the protein retinoblastoma [34, 35]. Furthermore, high ECM stiffness promotes this cyclin D1/Cdk4-dependent phosphorylation of retinoblastoma [4, 7, 19]. We showed that downregulation of survivin interferes with the stiffness-mediated induction of cyclin proteins, including cyclin D1 (expressed in early G1 phase), cyclin A (expressed in S phase), and cyclin B (expressed in G2/M phase) in VSMCs grown on stiff hydrogels. Furthermore, ectopic expression of survivin in VSMCs cultured on soft hydrogels was sufficient to promote cell cycling and proliferation. Together with previous studies, our findings suggest a novel mechanism by which the upregulation of survivin in VSMCs in response to ECM/arterial stiffness stimulates mechanosensitive cell cycling and proliferation.

The findings presented here expand on our work showing that stiffness-sensitive cell cycle progression and proliferation are associated with integrin, FAK, and Rac activity [4, 7, 19], suggesting that VSMCs sense ECM stiffness through integrin-dependent signaling. The inhibition of FAK or Rac interfered with stiffness-mediated expression of survivin. This likely involves FAK-Rac targeting of E2F1, which binds the survivin promotor [36] and induces survivin expression [37]. Furthermore, Rac1 binds and activates the transcription factor STAT3 [38], which also regulates the survivin promoter [39]. However, further studies are need to detail how FAK-Rac signaling triggers survivin transcription. Notably, the protein YAP also binds the survivin promoter and drives survivin transcription [40] and is involved in the mechanotransduction of ECM stiffness to cell cycling and proliferation [41].

In summary, we demonstrated that ECM/arterial stiffness signals an increase in survivin expression in VSMCs through FAK and Rac activation to induce cell cycle progression. This is a novel mechanism by which alterations to the microenvironment can trigger pathological phenotype switching of VSMCs from a contractile to a synthetic state. These findings also introduce potential targets for therapies for vascular diseases involving aberrant cell proliferation and arterial stiffness.

## METHODS

### Cell culture

Primary human VSMCs (hVSMCs; catalog number [cat. no.] 354-05a, Cell Applications, Inc.) were maintained in ≤90% Dulbecco’s modified Eagle’s medium (DMEM) supplemented with 1 mM sodium pyruvate, 2% MEM amino acids solution, 50 μg/mL gentamicin solution, 1% penicillin-streptomycin solution, and 10% fetal bovine serum (FBS). Primary mouse VSMCs (mVSMCs) were prepared from explant cultures of thoracic aortae from 2-month-old male C57BL/6 mice and maintained in ≤90% in low-glucose DMEM–F12 medium (1:1) supplemented with 10% FBS, 2 mM glutamine, 25 μM HEPES, 1%penicillin–streptomycin solution, and 50 μg/mL gentamicin solution. Both cell types were maintained in 10% CO_2_ at 37°C and used before passage 5. To synchronize VSMCs to the G0 cell cycle phase, cultures that were near confluence were incubated in serum-free DMEM containing 1 mg/ml heat-inactivated, fatty-acid-free bovine serum albumin (BSA) for 48 h. The starved cells were then treated with 0.05% trypsin–EDTA, centrifuged, resuspended, and plated on soft or stiff hydrogels with fresh medium containing 10%FBS for 24 h.

### siRNA transfection

hVSMCs were transfected with 200 nM survivin, FAK, or control siRNAs by using Lipofectamine 2000 reagent in Opti-MEM as previously described [7, 42]. After 4-5 h of siRNA transfection, cells were serum starved in fresh DMEM containing 1 mg/ml BSA for 43-44 h. All siRNA-based experiments were performed 72 h after transfection. The following survivin *(BIRC5)* and FAK *(PTK2)* siRNAs were obtained from Ambion: survivin siRNA #1 (ID no. 121294), 5’-CCACUUCCAGGGUUUAUUCtt-3’; survivin siRNA #2 (ID no. 121295), 5’-GCCAUUCUAAGUCAUUGGGtt-3’; FAK siRNA #1 (ID no. 157448), 5’-CCUAGCAGACUUUAACCAAtt-3’; and FAK siRNA #2 (ID no. 61352), 5’-GGCAUGGAGAUGCUACUGAtt-3’. A nonspecific siRNA (cat no. AM4636) was used as an experimental control.

### Adenovirus infection

hVMSCs were first incubated in DMEM containing 1 mg/ml BSA for 8-9 h. Cells were then incubated for 20-24 h with adenoviruses encoding wild-type survivin (cat. no. 1611, Vector Biolabs; multiplicity of infection [MOI], 25 and 50) or FRNK (a gift from the Assoian Laboratory; MOI, 600); adenoviruses encoding GFP (cat. no. 1060, Vector Biolabs; MOI, 50) or LacZ (a gift from the Assoian Laboratory; MOI, 600) were used as the respective experimental controls.

### Drug treatment

Serum-starved hVSMCs were plated on fibronectin-coated soft or stiff hydrogels with medium containing 10% FBS and were treated with 0.1, 0.5, or 2 μM YM155 (survivin inhibitor; cat. no. 11490, Cayman Chemical), 10 μM PF573228 (FAK inhibitor; cat. no. 14924, Cayman Chemical), 10 μM EHT1864 (Rac inhibitor; cat. no. 17258, Cayman Chemical), or dimethyl sulfoxide (vehicle control).

### Preparation of stiffness-tunable hydrogels

The soft (2–4 kPa) and stiff (16–30 kPa) polyacrylamide hydrogels [7, 43] approximate the physiological stiffness of a healthy mouse femoral artery and after vascular injury and atherosclerosis [4, 5], respectively. The protocol for generating stiffness-tunable polyacrylamide hydrogels was previously described [42, 43]. Briefly, glass coverslips were etched homogenously with a 0.1 M NaOH solution for 3 min and then treated with 3-(trimethoxysilyl)propyl methacrylate (cat. no. 440159, Sigma-Aldrich) to introduce amine groups to cross-link with the polyacrylamide hydrogel. Hydrogels of different stiffness were prepared by changing the ratio of 40% acrylamide to 1% bis-acrylamide in a mixed solution with sterilized water, ammonium persulfate (cat. no. A3678, Sigma-Aldrich), TEMED (cat. no. J63734.AC, Thermo Fisher Scientific), and an *N*-hydroxysuccinimide-fibronectin solution prepared by combining 1 part *N*-hydroxysuccinimide solution (cat. no. A8060, Sigma-Aldrich; 1 mg/ml in dimethyl sulfoxide) and 9 parts fibronectin solution (cat. no. 341631, Calbiochem; 100 μl fibronectin at 1 μg/μl dissolved in 1.9 ml Tris base [pH 8.4 pH]). Finally, the fibronectin-coated hydrogels were extensively washed in Dulbecco’s phosphate-buffered saline (DPBS) and water to remove unpolymerized polyacrylamide, and unreactive cross-linkers were blocked with 1 mg/ml BSA before VSMCs were seeded.

### RNA isolation and RT-qPCR

hVSMCs and mVSMCs cultured on soft or stiff hydrogels for 24 h were treated with TRIzol reagent to extract total RNA. The RNA was reverse transcribed and analyzed by RT-qPCR as previously described [42]. TaqMan probes (Invitrogen) were used for survivin (Mm00599749_m1 for mouse mRNA and Hs04194392_s1 for human mRNA), GAPDH (Mm99999915_g1 for mouse mRNA and Hs02786624_g1 for human mRNA), cyclin D1 (Hs00765553_m1), cyclin A (Hs00171105_m1), and cyclin B (Hs00259126_m1). The relative change in mRNA expression for each target mouse or human gene was determined by the comparative threshold cycle method using the gene for GAPDH as the reference.

### Protein extraction and immunoblotting

As previously described [42], total cell lysates were collected from hVSMCs or mVSMCs cultured on soft or stiff hydrogels by incubating the hydrogels face down for 2 min at room temperature on 5× sample buffer (250 mM Tris [pH 6.8], 10% SDS, 50% glycerol, 0.02% bromophenol blue, and 10 mM 2-mercaptoethanol). Equal amounts of extracted protein were fractionated on reducing 8–12% SDS-polyacrylamide gels, and the fractioned proteins were subsequently transferred electrophoretically onto polyvinylidene fluoride membranes via the Trans-Blot Turbo Transfer System (Bio-Rad). These membranes were blocked in 6% nonfat milk for 1.5 h at room temperature and then probed with antibodies against survivin (cat. no. NB500-201, Novus Biologicals; 1:200), cyclin D1 (cat. no. sc-20044, Santa Cruz Biotechnology; 1:200), cyclin A (a gift from the Assoian Laboratory (Kothapalli, 2003 #54), 1:500), FAK (cat. no. 39-6500, Invitrogen; 1:500), or GAPDH (cat. no. 60004-1-Ig or 10494-1-AP, Proteintech; 1:5,000). Antibody signals were detected using Clarity (cat. no. 1705061, Bio-Rad) or Clarity Max (cat. no. 1705062, Bio-Rad) Western ECL substrate.

### EdU incorporation

Serum-starved hVSMCs treated with control or survivin siRNAs or infected with adenovirus encoding wild-type survivin or GFP were plated on fibronectin-coated stiff or soft hydrogels with 10% FBS and then incubated with 20 μM EdU for 24 or 36 h. Cells were fixed in 3.7% formaldehyde and visualized using the Click-iT EdU Alexa Fluor 594 imaging kit (cat. no. C10339, Invitrogen) according to the manufacturer’s instructions. Nuclei were stained with DAPI (4’,6-diamidino-2-phenylindole), and coverslips were mounted on microscope slides. Three to eight fields of view were counted per coverslip to determine the percentage of hVSMCs with DAPI-stained nuclei that were positive for EdU.

### Bioinformatics analysis

#### (i) Gene expression analysis

Differential gene expression analysis was performed on raw microarray data. Duplicate and blank (no name) gene entries were removed from both data lists, and genes with nonsignificant differential expression values were filtered out before further analysis. For *in vitro* microarray data, differentially expressed genes (DEGs) were defined as genes having a ≥1.5-fold change and a false-discovery rate (FDR)-adjusted *p* value *(q* value) of ≤0.05. For the *in vivo* microarray data, DEGs were defined as genes having a ≥2.0-fold change and a *q* value of ≤0.15. Python’s bioinfokit package was used to generate volcano plots and heat maps with hierarchical clustering of DEGs.

#### (ii) Functional enrichment analysis

Functional enrichment analysis was performed using the g:GOSt tool in gProfiler (https://biit.cs.ut.ee/gprofiler/gost). The statistical domain scope of the analysis was only annotated genes, and the significance threshold was set to the g:SCS algorithm for computing multiple-testing correction for *p* values acquired from Gene Ontology (GO). Significant GO terms were defined by an adjusted *p* value of ≤0.05. The top 25 Biological Process GO terms were presented in histograms on a scale of −log(adjusted *p* value). Cytoscape (https://cytoscape.org/) was used to visualize the functional enrichment results as they pertained to DEGs common to both the *in vitro* and *in vivo* data sets. The list of 74 commonly upregulated DEGs was input into Cytoscape’s String application as a protein query, and the *in vitro* expression data were imported into the node table so that the log_2_(fold change) values could be used to indicate expression levels, with node color indicating intensity. The only nodes kept were those in the network that displayed a relationship to any other gene in the network at a score of 0.95 and fell under the following Biological Process GO categories: cell cycle (GO:0007049), regulation of cell cycle (GO:0051726), mitotic cell cycle (GO:0000278), mitotic cell cycle processes (GO:1903047), G2/M transition of mitotic cell cycle (GO:0000086), cell division (GO:0051301), nuclear division (GO:0000280), and mitotic nuclear division (GO:0140014). The resulting network contained 37 genes.

#### (iii) Network analysis

Ingenuity Pathway Analysis (IPA, Qiagen) was used to perform further bioinformatics analysis on the filtered microarray data. A core analysis was run on each of the data sets, which returned information on various mechanistic pathways and enriched functions based on the literature compiled in the Ingenuity Knowledge Base. The “Diseases and Functions” tool was used to identify molecules known to be involved in cell cycle progression within the *in vitro* data set, and the “My Pathway” tool was subsequently used to display known relationships between *Birc5* and other genes within the cell cycle progression function. The Z-directional components of the expression analysis were based on the expression log ratio values. Functions with a Z-score of >2 were regarded as having significant activation, whereas those with a Z-score of <-2 were considered as having significant inhibition. The “Molecule Activity Predictor” tool was used to display gene expression levels via node color and intensity and to generate predicted activation states of molecules and interactions based on the results of the Core Analysis.

### Statistical analysis

Statistical significance was determined by paired, two-tailed Student’s *t* tests. Graphs show means + SEMs from the indicated number of independent experiments.

## ACKNOWLEDGEMENTS

This work was supported by American Heart Association Career Development Award (18CDA34080415 to Y.B.) and NIH/NHLBI grant 1R56HL163168-01 (to Y.B.).

## AUTHOR CONTRIBUTIONS

J.C.B., A.S., S-J.H, J.K., K.L., and Y.B. designed the research. J.C.B., A.S., J.A.B., A.K., Y.H., and Y.B. performed cell/molecular biology experiments and data analysis. A.S., K.E.P., V.M.T., and Y.B. performed bioinformatic analysis. J.C.B. and Y.B. performed statistical tests. J.C.B., A.S., J.A.B., S-J.H, J.K., K.L., and Y.B. wrote and edited the manuscript.

## SUPPLEMENTARY FIGURE LEGENDS

**Figure S1.**
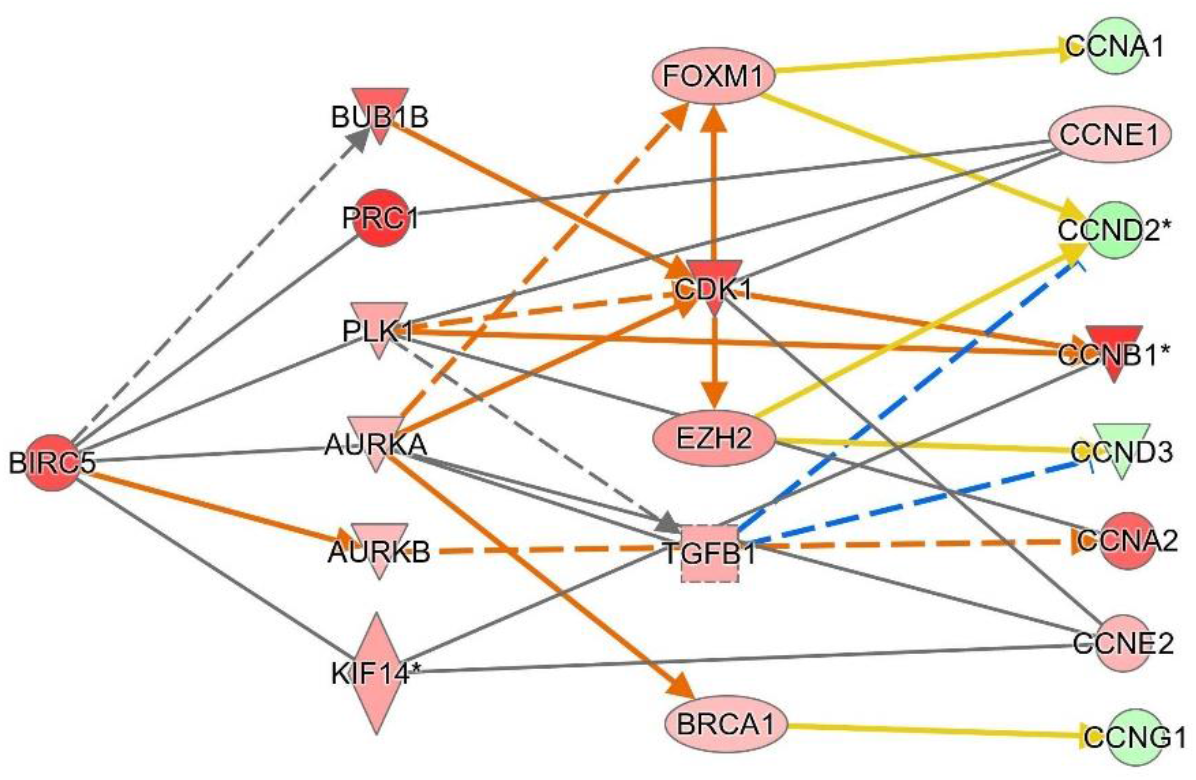
Predictive model for survivin (BIRC5) regulation of cyclins. Network diagram displaying downstream targets of BIRC5 and potential intermediate molecules, with node color and intensity representing the observed expression levels or predicted activation states based on the *in vivo* data.

**Figure S2.**
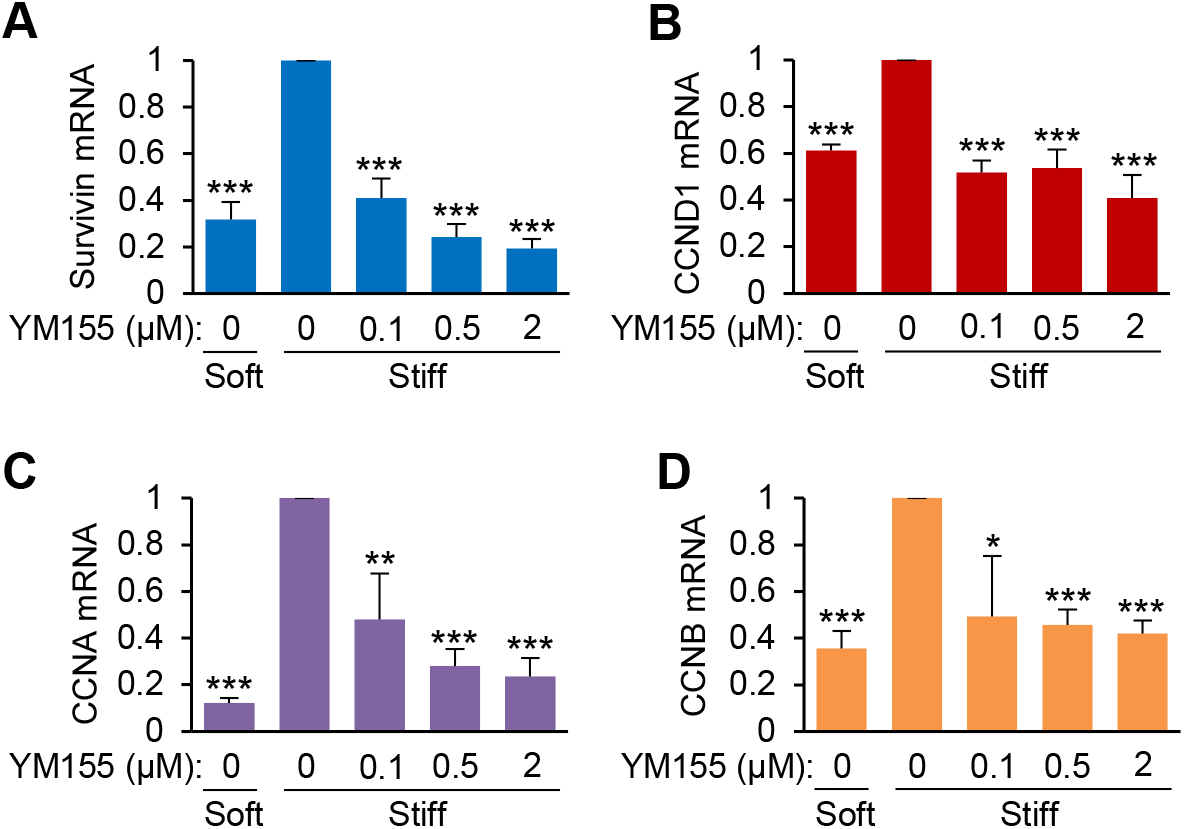
Pharmacologic reduction of survivin expression decreases cyclin D1, A, and B mRNA expression in human vascular smooth muscle cells (hVSMCs). G0-synchronized hVSMCs were seeded on soft or stiff hydrogels with or without YM155 (inhibits survivin expression) for 24 h. RT-qPCR was performed to assess the expression of survivin (**A**), cyclin D1 (CCND1; **B**), cyclin A (CCNA; **C**), and cyclin B (CCNB; **D**) mRNA; levels were normalized to those in hVSMCs treated with dimethyl sulfoxide (0 μM YM155) on stiff hydrogels. *n* = 10–14 (A), *n* = 3–5 (B), *n* = 5–9 (C), and *n* = 3–7 (D) independent experiments. Data are means + SEMs. *\*p* < 0.05, \*\**p* < 0.01, \*\*\**p* < 0.001.

## TABLES

**Table S1.** Differentially expressed gene (DEG) lists of the *in vivo* study

**Table S2.** Differentially expressed gene (DEG) lists of the *in vitro* study

## BIBLIOGRAPHY

1. Mitchell, G.F., Hwang, S.-J., Vasan, R.S., Larson, M.G., Pencina, M.J., Hamburg, N.M., Vita, J.A., Levy, D., and Benjamin, E.J. (2010). Arterial Stiffness and Cardiovascular Events: The Framingham Heart Study. Circulation 121, 505–511.

2. Liao, D., Arnett, D.K., Tyroler, H.A., Riley, W.A., Chambless, L.E., Szklo, M., and Heiss, G. (1999). Arterial Stiffness and the Development of Hypertension. Hypertension 34, 201.

3. Mattace-Raso, F.U.S., van der Cammen, T.J.M., Hofman, A., van Popele, N.M., Bos, M.L., Schalekamp, M.A.D.H., Asmar, R., Reneman, R.S., Hoeks, A.P.G., Breteler, M.M.B., et al. (2006). Arterial Stiffness and Risk of Coronary Heart Disease and Stroke: The Rotterdam Study. Circulation 113, 657–663.

4. Klein, E.A., Yin, L., Kothapalli, D., Castagnino, P., Byfield, F.J., Xu, T., Levental, I., Hawthorne, E., Janmey, P.A., and Assoian, R.K. (2009). Cell-Cycle Control by Physiological Matrix Elasticity and In Vivo Tissue Stiffening. Current Biology 19, 1511–1518.

5. Kothapalli, D., Liu, S.-L., Bae, Yong H., Monslow, J., Xu, T., Hawthorne, Elizabeth A., Byfield, Fitzroy J., Castagnino, P., Rao, S., Rader, Daniel J., et al. (2012). Cardiovascular Protection by ApoE and ApoE-HDL Linked to Suppression of ECM Gene Expression and Arterial Stiffening. Cell Reports 2, 1259–1271.

6. Hahn, C., and Schwartz, M.A. (2009). Mechanotransduction in vascular physiology and atherogenesis. Nature reviews. Molecular cell biology 10, 53–62.

7. Bae, Y.H., Mui, K.L., Hsu, B.Y., Liu, S.L., Cretu, A., Razinia, Z., Xu, T., Pure, E., and Assoian, R.K. (2014). A FAK-Cas-Rac-lamellipodin signaling module transduces extracellular matrix stiffness into mechanosensitive cell cycling. Science signaling 7, ra57.

8. Thyberg, J., Hedin, U., Sjölund, M., Palmberg, L., and Bottger, B.A. (1990). Regulation of differentiated properties and proliferation of arterial smooth muscle cells. Arteriosclerosis, Thrombosis, and Vascular Biology 10, 966–990.

9. Owens, G.K., Kumar, M.S., and Wamhoff, B.R. (2004). Molecular Regulation of Vascular Smooth Muscle Cell Differentiation in Development and Disease, Volume 84.

10. Owens, G.K. (1995). Regulation of differentiation of vascular smooth muscle cells, Volume 75.

11. Thyberg, J., Blomgren, K., Roy, J., Tran, P.K., and Hedin, U. (1997). Phenotypic Modulation of Smooth Muscle Cells after Arterial Injury Is Associated with Changes in the Distribution of Laminin and Fibronectin. Journal of Histochemistry & Cytochemistry 45, 837–846.

12. Liu, S.-L., Bae, Y.H., Yu, C., Monslow, J., Hawthorne, E.A., Castagnino, P., Branchetti, E., Ferrari, G., Damrauer, S.M., Puré, E., et al. (2015). Matrix metalloproteinase-12 is an essential mediator of acute and chronic arterial stiffening. Sci Rep 5, 17189.

13. von Kleeck, R., Roberts, E., Castagnino, P., Bruun, K., Brankovic, S.A., Hawthorne, E.A., Xu, T., Tobias, J.W., and Assoian, R.K. (2021). Arterial stiffness and cardiac dysfunction in Hutchinson-Gilford Progeria Syndrome corrected by inhibition of lysyl oxidase. Life science alliance 4.

14. Davies, P.F. (2009). Hemodynamic shear stress and the endothelium in cardiovascular pathophysiology. Nat Clin Pract Cardiovasc Med 6, 16–26.

15. Kaess, B.M., Rong, J., Larson, M.G., Hamburg, N.M., Vita, J.A., Levy, D., Benjamin, E.J., Vasan, R.S., and Mitchell, G.F. (2012). Aortic stiffness, blood pressure progression, and incident hypertension. Jama 308, 875–881.

16. Boutouyrie, P., Chowienczyk, P., Humphrey, J.D., and Mitchell, G.F. (2021). Arterial Stiffness and Cardiovascular Risk in Hypertension. Circulation research 128, 864–886.

17. Ikonomidis, I., Makavos, G., and Lekakis, J. (2015). Arterial stiffness and coronary artery disease. Current opinion in cardiology 30, 422–431.

18. Huang, X., Yang, N., Fiore, V.F., Barker, T.H., Sun, Y., Morris, S.W., Ding, Q., Thannickal, V. J., and Zhou, Y. (2012). Matrix stiffness-induced myofibroblast differentiation is mediated by intrinsic mechanotransduction. American journal of respiratory cell and molecular biology 47, 340–348.

19. Mui, Keeley L., Bae, Yong H., Gao, L., Liu, S.-L., Xu, T., Radice, Glenn L., Chen, Christopher S., and Assoian, Richard K. (2015). N-Cadherin Induction by ECM Stiffness and FAK Overrides the Spreading Requirement for Proliferation of Vascular Smooth Muscle Cells. Cell Reports 10, 1477–1486.

20. Razinia, Z., Castagnino, P., Xu, T., Vázquez-Salgado, A., Puré, E., and Assoian, R.K. (2017). Stiffness-dependent motility and proliferation uncoupled by deletion of CD44. Sci Rep 7, 16499–16499.

21. Liu, F., Mih, J.D., Shea, B.S., Kho, A.T., Sharif, A.S., Tager, A.M., and Tschumperlin, D.J. (2010). Feedback amplification of fibrosis through matrix stiffening and COX-2 suppression. The Journal of Cell Biology 190, 693–706.

22. Peyton, S.R., and Putnam, A.J. (2005). Extracellular matrix rigidity governs smooth muscle cell motility in a biphasic fashion. Journal of Cellular Physiology 204, 198–209.

23. Yeung, T., Georges, P.C., Flanagan, L.A., Marg, B., Ortiz, M., Funaki, M., Zahir, N., Ming, W., Weaver, V., and Janmey, P.A. (2005). Effects of substrate stiffness on cell morphology, cytoskeletal structure, and adhesion. Cell Motility and the Cytoskeleton 60, 24–34.

24. Solon, J., Levental, I., Sengupta, K., Georges, P.C., and Janmey, P.A. (2007). Fibroblast Adaptation and Stiffness Matching to Soft Elastic Substrates. Biophysical Journal 93, 4453–4461.

25. Blanc-Brude, O.P., Yu, J., Simosa, H., Conte, M.S., Sessa, W.C., and Altieri, D.C. (2002). Inhibitor of apoptosis protein survivin regulates vascular injury. Nat Med 8, 987–994.

26. Simosa, H.F., Wang, G., Sui, X., Peterson, T., Narra, V., Altieri, D.C., and Conte, M.S. (2005). Survivin expression is up-regulated in vascular injury and identifies a distinct cellular phenotype. Journal of Vascular Surgery 41, 682–690.

27. Hoel, A.W., Yu, P., Nguyen, K.P., Sui, X., Plescia, J., Altieri, D.C., and Conte, M.S. (2012). Mitochondrial Heat Shock Protein-90 Modulates Vascular Smooth Muscle Cell Survival and the Vascular Injury Response in Vivo. The American Journal of Pathology 181, 1151–1157.

28. Ambrosini, G., Adida, C., and Altieri, D.C. (1997). A novel anti-apoptosis gene, survivin, expressed in cancer and lymphoma. Nat Med 3, 917–921.

29. Mehrotra, S., Languino, L.R., Raskett, C.M., Mercurio, A.M., Dohi, T., and Altieri, D.C. IAP Regulation of Metastasis. Cancer Cell 17, 53–64.

30. Kobayashi, K., Hatano, M., Otaki, M., Ogasawara, T., and Tokuhisa, T. (1999). Expression of a murine homologue of the inhibitor of apoptosis protein is related to cell proliferation. Proceedings of the National Academy of Sciences 96, 1457–1462.

31. Horowitz, J.C., Ajayi, I.O., Kulasekaran, P., Rogers, D.S., White, J.B., Townsend, S.K., White, E.S., Nho, R.S., Higgins, P.D., Huang, S.K., et al. (2012). Survivin expression induced by endothelin-1 promotes myofibroblast resistance to apoptosis. The international journal of biochemistry & cell biology 44, 158–169.

32. Yoshida, T., Zhang, Y., Rivera Rosado, L.A., Chen, J., Khan, T., Moon, S.Y., and Zhang, B. (2010). Blockade of Rac1 activity induces G1 cell cycle arrest or apoptosis in breast cancer cells through downregulation of cyclin D1, survivin, and X-linked inhibitor of apoptosis protein. Molecular cancer therapeutics 9, 1657–1668.

33. Krämer, A., Green, J., Pollard, J., Jr., and Tugendreich, S. (2014). Causal analysis approaches in Ingenuity Pathway Analysis. Bioinformatics (Oxford, England) 30, 523–530.

34. Ito, T., Shiraki, K., Sugimoto, K., Yamanaka, T., Fujikawa, K., Ito, M., Takase, K., Moriyama, M., Kawano, H., Hayashida, M., et al. (2000). Survivin promotes cell proliferation in human hepatocellular carcinoma. Hepatology (Baltimore, Md.) 31, 1080–1085.

35. Connell, C.M., Wheatley, S.P., and McNeish, I.A. (2008). Nuclear survivin abrogates multiple cell cycle checkpoints and enhances viral oncolysis. Cancer research 68, 7923–7931.

36. Liu, L., Zhang, H., Shi, L., Zhang, W., Yuan, J., Chen, X., Liu, J., Zhang, Y., and Wang, Z. (2014). Inhibition of Rac1 activity induces G1/S phase arrest through the GSK3/cyclin D1 pathway in human cancer cells. Oncology reports 32, 1395–1400.

37. Jiang, Y., Saavedra, H.I., Holloway, M.P., Leone, G., and Altura, R.A. (2004). Aberrant regulation of survivin by the RB/E2F family of proteins. The Journal of biological chemistry 279, 40511–40520.

38. Simon, A.R., Vikis, H.G., Stewart, S., Fanburg, B.L., Cochran, B.H., and Guan, K.L. (2000). Regulation of STAT3 by direct binding to the Rac1 GTPase. Science (New York, N.Y.) 290, 144–147.

39. Gritsko, T., Williams, A., Turkson, J., Kaneko, S., Bowman, T., Huang, M., Nam, S., Eweis, I., Diaz, N., Sullivan, D., et al. (2006). Persistent activation of stat3 signaling induces survivin gene expression and confers resistance to apoptosis in human breast cancer cells. Clinical cancer research : an official journal of the American Association for Cancer Research 12, 11–19.

40. Rosenbluh, J., Nijhawan, D., Cox, A.G., Li, X., Neal, J.T., Schafer, E.J., Zack, T.I., Wang, X., Tsherniak, A., Schinzel, A.C., et al. (2012). β-Catenin-driven cancers require a YAP1 transcriptional complex for survival and tumorigenesis. Cell 151, 1457–1473.

41. Dupont, S., Morsut, L., Aragona, M., Enzo, E., Giulitti, S., Cordenonsi, M., Zanconato, F., Le Digabel, J., Forcato, M., Bicciato, S., et al. (2011). Role of YAP/TAZ in mechanotransduction. Nature 474, 179–183.

42. Brazzo, J.A., Biber, J.C., Nimmer, E., Heo, Y., Ying, L., Zhao, R., Lee, K., Krause, M., and Bae, Y. (2021). Mechanosensitive expression of lamellipodin promotes intracellular stiffness, cyclin expression and cell proliferation. Journal of cell science 134.

43. Cretu, A., Castagnino, P., and Assoian, R. (2010). Studying the Effects of Matrix Stiffness on Cellular Function using Acrylamide-based Hydrogels. e2089.

