## Supplementary material for "Survivin is a mechanosensitive cell cycle regulator in vascular smooth muscle cells": Table 1

**Table S1. Differentially expressed gene (DEG) lists of the *in vivo* study**

| Gene Symbol | p-value(inj vs. control) | q-value(inj vs control) | Fold-Change(inj vs. control) | log2FoldChange |
|---|---|---|---|---|
| 0610010O12A4:A652Rik | 0.00724946 | 0.129666997 | -2.02767 | -1.01981965 |
| 1110067D22Rik | 0.00966545 | 0.135920391 | -2.19834 | -1.136416718 |
| 1300014I06Rik | 0.00625995 | 0.125597723 | -2.0226 | -1.016211417 |
| 1700019E19Rik | 0.00131066 | 0.105261221 | -2.19917 | -1.136959153 |
| 2010002N04Rik | 0.00646528 | 0.126320254 | 3.65658 | 1.870494926 |
| 2310030G06Rik | 0.00153563 | 0.108027569 | -2.36306 | -1.240660059 |
| 2700078K21Rik | 0.00880796 | 0.134106158 | -2.64516 | -1.403354857 |
| 2810417H13Rik | 0.000457496 | 0.109915353 | 6.30295 | 2.656027218 |
| 2810417H13Rik | 0.00182907 | 0.106915741 | 2.62759 | 1.39374018 |
| 2810432L12Rik | 0.00273717 | 0.110697246 | -2.18548 | -1.12794824 |
| 4632434I11Rik | 0.0101027 | 0.137631689 | 2.35949 | 1.238475057 |
| 4930506M07Rik | 0.0107023 | 0.139820371 | 2.56556 | 1.359273766 |
| 4930562F07Rik | 0.00978766 | 0.136622433 | -2.08472 | -1.059855806 |
| 4933439F18Rik | 0.0121168 | 0.145617329 | -2.01794 | -1.012879997 |
| 5730407I07Rik | 0.000799308 | 0.097674059 | -2.27502 | -1.185877834 |
| 5830454E08Rik | 0.00800818 | 0.131005137 | -2.46663 | -1.302539302 |
| 6330403A02Rik | 0.000839542 | 0.097946567 | -6.24187 | -2.641981904 |
| A430107O13Rik | 0.00223408 | 0.108825031 | -2.22373 | -1.152981252 |
| A830039N20Rik | 0.00116521 | 0.102589141 | -3.22348 | -1.688617792 |
| AA467197 | 0.00311912 | 0.112217071 | 14.6613 | 3.873941126 |
| Abcc3 | 0.0110179 | 0.141019172 | 2.4092 | 1.268554164 |
| Abra | 0.000385914 | 0.109406619 | -9.63339 | -3.268038261 |
| Acsl1 | 0.00403316 | 0.114454541 | -2.69174 | -1.4285387 |
| Acss1 | 0.00304221 | 0.111863364 | -2.28063 | -1.189433506 |
| Actc1 | 0.00204789 | 0.107514225 | -7.68216 | -2.941508937 |
| Actg2 | 0.00600844 | 0.124882166 | -4.36645 | -2.126460802 |
| Actn1 | 0.000371566 | 0.107488736 | -2.66561 | -1.414468228 |
| Actn1 | 0.00465141 | 0.115978429 | -2.7597 | -1.464512349 |
| Actn4 | 0.00820015 | 0.131937714 | -2.30499 | -1.204761694 |
| Adam12 | 0.00714886 | 0.129171562 | 2.99087 | 1.580565204 |
| Adam19 | 0.00207509 | 0.107351828 | 2.29157 | 1.196336356 |
| Adamts8 | 0.00209906 | 0.108000637 | -4.33782 | -2.116973587 |
| Adamtsl1 | 0.00187965 | 0.107219472 | -2.4099 | -1.268973204 |
| Adcy5 | 0.0114259 | 0.14282375 | -4.94299 | -2.305381856 |
| Adh7 | 0.00643876 | 0.126411943 | -4.27707 | -2.096622312 |
| AF251705 | 0.00194854 | 0.107682474 | 4.58993 | 2.198472152 |
| Agtr1b | 0.0114061 | 0.142891266 | -3.33995 | -1.739824962 |
| Ahr | 0.00135786 | 0.106047744 | 2.60665 | 1.382196882 |
| Aif1 | 0.0114596 | 0.143118793 | 7.58425 | 2.923006521 |
| Ak4 | 0.0105347 | 0.138975684 | -2.48844 | -1.315238702 |
| Ak4 | 0.00497011 | 0.118704818 | -2.85572 | -1.513856126 |
| Akap12 | 0.0106311 | 0.13921094 | -2.30081 | -1.202143655 |
| Akap6 | 0.00264701 | 0.11117442 | -3.83748 | -1.940162981 |
| Akr1b8 | 0.00287392 | 0.1108512 | 2.24381 | 1.165950517 |
| Akr1c14 | 0.000756281 | 0.095716814 | -2.22865 | -1.15617048 |
| Alcam | 0.00385843 | 0.114063077 | 4.10487 | 2.037336533 |
| Amigo2 | 0.00321635 | 0.112019069 | -5.48945 | -2.45665854 |
| Ammecr1 | 0.00321262 | 0.112303054 | 2.13866 | 1.096707144 |
| Amy1 | 0.0126927 | 0.14729351 | -3.57391 | -1.837501196 |

| | | | | |
|---|---|---|---|---|
| Angpt2 | 0.00519408 | 0.12089669 | 2.00489 | 1.003523084 |
| Anln | 0.0022214 | 0.1085805 | 3.37733 | 1.755883152 |
| Anpep | 0.00112094 | 0.10152923 | 2.17008 | 1.117748229 |
| Anxa8 | 0.00714199 | 0.129294647 | 5.33495 | 2.41547475 |
| Ap1s2 | 0.00028398 | 0.113392014 | 2.36446 | 1.241510736 |
| Apbb1ip | 0.00654727 | 0.126526997 | 2.49075 | 1.316580224 |
| Apcdd1 | 0.00425801 | 0.115737856 | -2.86578 | -1.518928435 |
| Aplnr | 0.00476902 | 0.117057764 | 12.4067 | 3.633047526 |
| Apobec1 | 0.00619985 | 0.125636703 | 4.47921 | 2.163244306 |
| Apoe | 0.00266931 | 0.110635875 | 2.78421 | 1.477268031 |
| Arhgap11a | 0.00877647 | 0.133914383 | 3.25781 | 1.703902466 |
| Arhgap25 | 0.00129895 | 0.10521495 | 2.36483 | 1.241736477 |
| Arhgap30 | 0.00530621 | 0.121315366 | 2.8751 | 1.523612136 |
| Arhgef9 | 0.000246478 | 0.118434768 | -2.7786 | -1.474356044 |
| Arrb2 | 0.00907272 | 0.13445458 | 3.66106 | 1.872261418 |
| Aspm | 0.00296644 | 0.11109455 | 2.95843 | 1.56483176 |
| Atf3 | 0.00965659 | 0.135930649 | -3.22736 | -1.690358127 |
| Atp1a2 | 0.010032 | 0.13732844 | -2.75432 | -1.461696262 |
| Atp2a3 | 0.0011994 | 0.103352553 | -2.34158 | -1.22748594 |
| B230118H07Rik | 0.010564 | 0.138845341 | -2.1061 | -1.074571698 |
| B3galt1 | 0.000309566 | 0.104478525 | -2.84972 | -1.510822922 |
| Baz1a | 0.0129357 | 0.148052925 | 3.49804 | 1.806546787 |
| Bcar3 | 0.0100743 | 0.137509102 | -2.01887 | -1.013549744 |
| Bcl2a1a | 0.00453005 | 0.11590877 | 17.4214 | 4.12278866 |
| Bcl2a1b | 0.00457339 | 0.115867387 | 16.2684 | 4.024000464 |
| Bcl2a1c | 0.0122331 | 0.145779061 | 4.27385 | 2.095536274 |
| Bcl2a1d | 0.00315359 | 0.11231693 | 14.7954 | 3.887076796 |
| Bcl2l11 | 0.00484712 | 0.117549916 | 2.01873 | 1.013447967 |
| Bicd1 | 0.00306471 | 0.112109069 | -2.30355 | -1.203860792 |
| Birc5 | 0.00862726 | 0.133142526 | 3.98644 | 1.995100955 |
| Bmper | 0.00345885 | 0.112452291 | 3.74936 | 1.906644355 |
| Bnc2 | 0.00300945 | 0.111965758 | 2.10772 | 1.075683225 |
| Bub1 | 0.0011917 | 0.10331711 | 5.78986 | 2.533528464 |
| Bub1b | 0.000101914 | 0.103187925 | 3.58758 | 1.843011003 |
| C130021O09Rik | 0.00330404 | 0.111246477 | -2.18174 | -1.125478421 |
| C1qa | 0.00175652 | 0.108020265 | 5.95639 | 2.574438219 |
| C1qb | 0.00950986 | 0.135479664 | 3.34184 | 1.740642662 |
| C3ar1 | 0.00531602 | 0.121343935 | 6.03911 | 2.594335951 |
| C5ar1 | 0.00617599 | 0.125602092 | 2.89889 | 1.535500591 |
| C79407 | 0.0025247 | 0.111661849 | 2.76684 | 1.468239219 |
| Cabp1 | 0.00777454 | 0.130728475 | -2.14395 | -1.100267512 |
| Cacnb2 | 0.00732684 | 0.130147816 | -3.91183 | -1.96784272 |
| Cald1 | 0.00206296 | 0.107115231 | -2.13667 | -1.095360995 |
| Camk2g | 0.00767888 | 0.130670017 | -3.27528 | -1.711620174 |
| Cap2 | 0.00955272 | 0.135342135 | -5.70058 | -2.511106629 |
| Capg | 0.00228041 | 0.109205445 | 2.26714 | 1.180873483 |
| Capn6 | 0.00795817 | 0.131094782 | 2.32364 | 1.21638657 |
| Car9 | 0.00502774 | 0.119177616 | -2.36079 | -1.239266376 |
| Casc5 | 0.00052614 | 0.105042739 | 6.66524 | 2.736656824 |
| Casp3 | 0.00463445 | 0.116168574 | 2.44316 | 1.288748347 |
| Ccdc107 | 0.00930309 | 0.135252616 | -2.22553 | -1.154146285 |
| Ccdc109b | 0.0128322 | 0.14788328 | 2.46711 | 1.302822042 |

| | | | | |
|---|---|---|---|---|
| Ccdc3 | 0.0124668 | 0.146410017 | -3.22219 | -1.688041246 |
| Ccna2 | 0.000364623 | 0.10767773 | 3.50623 | 1.809920636 |
| Ccnb1 | 0.00812369 | 0.131453545 | 4.73298 | 2.242748826 |
| Ccnb1 | 0.00758575 | 0.130416017 | 4.47735 | 2.1626451 |
| Ccnb1 | 0.0070238 | 0.129385789 | 3.79277 | 1.923251887 |
| Ccr2 | 0.0102134 | 0.13801234 | 8.64685 | 3.112174663 |
| Ccr5 | 0.00778374 | 0.130689386 | 11.3239 | 3.501299009 |
| Ccr5 | 0.00778374 | 0.130689386 | 11.3239 | 3.501299009 |
| Cd109 | 0.00174689 | 0.10766159 | 2.59214 | 1.374143639 |
| Cd163 | 0.00739493 | 0.129892358 | -4.40317 | -2.138543216 |
| Cd209f | 0.00437189 | 0.115403242 | -2.79178 | -1.481186924 |
| Cd38 | 0.000855809 | 0.097048741 | 2.63094 | 1.395578347 |
| Cd3g | 0.0131676 | 0.149499984 | 9.63418 | 3.268161879 |
| Cd48 | 0.00850934 | 0.133134541 | 6.25556 | 2.645139041 |
| Cd52 | 0.0115386 | 0.143725532 | 13.5336 | 3.758473749 |
| Cd53 | 0.00575181 | 0.123533192 | 7.71897 | 2.948408351 |
| Cd59b | 0.00354399 | 0.112763318 | -2.31928 | -1.213677985 |
| Cd68 | 0.00708882 | 0.129489721 | 5.28031 | 2.400622631 |
| Cd72 | 0.00419323 | 0.115191929 | 11.3667 | 3.506741564 |
| Cd80 | 0.00414578 | 0.114778187 | 2.70011 | 1.433018183 |
| Cd84 | 0.00275517 | 0.110012774 | 5.04652 | 2.335288871 |
| Cdc25c | 0.00351342 | 0.112931357 | 2.23431 | 1.159829367 |
| Cdc42ep3 | 0.00106036 | 0.101558128 | -2.5433 | -1.346705134 |
| Cdca8 | 0.00190448 | 0.106914867 | 2.36638 | 1.242681764 |
| Cdh11 | 0.00525336 | 0.12098518 | 2.5823 | 1.368656616 |
| Cdh19 | 0.00498054 | 0.118753834 | -2.80325 | -1.487099592 |
| Cdk1 | 0.000627629 | 0.09723105 | 4.13613 | 2.048281531 |
| Cdk14 | 0.00378585 | 0.114058286 | 2.05783 | 1.041123804 |
| Cdkl5 | 0.0000767 | 0.098838409 | -4.66947 | -2.223259261 |
| Cenpe | 0.000534593 | 0.103100079 | 5.68252 | 2.506530857 |
| Cenpf | 0.00680569 | 0.127945167 | 3.59694 | 1.846770094 |
| Cenph | 0.00618547 | 0.125524749 | 3.34996 | 1.744143869 |
| Cenpk | 0.00434987 | 0.115683691 | 4.46589 | 2.158947716 |
| Cfi | 0.00150147 | 0.107220843 | 4.03028 | 2.010880072 |
| Chd7 | 0.0121649 | 0.145701274 | 4.70795 | 2.235098999 |
| Chd7 | 0.0107239 | 0.140038031 | 4.45673 | 2.155985561 |
| Chd7 | 0.00512801 | 0.120446631 | 4.45339 | 2.15490396 |
| Chd7 | 0.00837732 | 0.132754065 | 3.73068 | 1.899438618 |
| Chd7 | 0.00461159 | 0.115800333 | 2.91936 | 1.545652127 |
| Chek2 | 0.000169472 | 0.123193108 | 2.05887 | 1.041852739 |
| Chmp4c | 0.00292035 | 0.110981129 | -2.81106 | -1.491113003 |
| Chn2 | 0.00343502 | 0.112192185 | -2.16791 | -1.116304119 |
| Chpt1 | 0.00238814 | 0.110266725 | -2.4861 | -1.313885887 |
| Ckb | 0.000567729 | 0.101227152 | -2.37068 | -1.245300595 |
| Cklf | 0.00559718 | 0.122912512 | 3.00093 | 1.585409667 |
| Cks2 | 0.0048016 | 0.116946186 | 7.87041 | 2.976438793 |
| Cks2 | 0.00477532 | 0.117111005 | 7.28555 | 2.865037888 |
| Cks2 | 0.00428293 | 0.11563911 | 5.66626 | 2.502396802 |
| Clec12a | 0.0120506 | 0.145562211 | 8.50206 | 3.08781244 |
| Clec4d | 0.00433815 | 0.115588865 | 9.29881 | 3.217046101 |
| Clec4n | 0.000247672 | 0.115106577 | 12.6665 | 3.66294603 |
| Clec7a | 0.00221246 | 0.108894516 | 4.1575 | 2.055716264 |

| | | | | |
|---|---|---|---|---|
| Clybl | 0.00204483 | 0.107953316 | -2.36303 | -1.240639604 |
| Cmbl | 0.00303016 | 0.111855516 | -2.2507 | -1.170371224 |
| Cmtm3 | 0.00276251 | 0.109687897 | 2.00741 | 1.005335308 |
| Col14a1 | 0.00110313 | 0.101868845 | 2.91964 | 1.545790492 |
| Col19a1 | 0.000201429 | 0.118969003 | -4.23036 | -2.080777398 |
| Col1a2 | 0.00852211 | 0.133187331 | 2.39269 | 1.258633492 |
| Col3a1 | 0.0111416 | 0.14196151 | 2.524 | 1.33571191 |
| Col5a2 | 0.00155928 | 0.107035322 | 2.36337 | 1.24084551 |
| Comp | 0.00348043 | 0.113024273 | 2.68422 | 1.42450292 |
| Coro1a | 0.00510457 | 0.120394808 | 4.10282 | 2.036615861 |
| Cox7a1 | 0.00773368 | 0.13050585 | -2.04297 | -1.030663445 |
| Cryab | 0.00085462 | 0.097695472 | -2.11765 | -1.082465894 |
| Csrp1 | 0.00822331 | 0.131786794 | -3.67451 | -1.877552565 |
| Cthrc1 | 0.000474572 | 0.109383059 | 6.90264 | 2.787148244 |
| Ctnna3 | 0.0126489 | 0.147388539 | -5.46242 | -2.449540582 |
| Ctsb | 0.00139164 | 0.106629714 | 2.16643 | 1.115319622 |
| Ctsc | 0.00606448 | 0.12494768 | 3.36825 | 1.751999223 |
| Ctsk | 0.00358112 | 0.112430512 | 2.8265 | 1.499016697 |
| Ctss | 0.00313466 | 0.112348434 | 7.76467 | 2.956924611 |
| Cx3cr1 | 0.00788397 | 0.130937639 | 3.53067 | 1.819941983 |
| Cxcl12 | 0.000408321 | 0.109206607 | 4.36456 | 2.12583622 |
| Cxcl16 | 0.0119686 | 0.145376954 | 4.18154 | 2.064034364 |
| Cxcr4 | 0.0000125 | 0.050625 | 6.97132 | 2.801431852 |
| Cyba | 0.00776772 | 0.13084662 | 2.66026 | 1.411567254 |
| Cybb | 0.00380106 | 0.113911259 | 9.33136 | 3.222087362 |
| Cyp7b1 | 0.00690223 | 0.128482088 | 3.51438 | 1.813270194 |
| Cys1 | 0.0000867 | 0.102414375 | -2.34392 | -1.228922388 |
| Cyth4 | 0.00799366 | 0.131145984 | 2.80467 | 1.487831032 |
| Cytl1 | 0.00220991 | 0.109147994 | -16.7066 | -4.062345857 |
| D17H6S56E-5 | 0.0016961 | 0.107331328 | 3.46251 | 1.791818238 |
| D2Ertd750e | 0.00181248 | 0.107497506 | 2.19762 | 1.135941945 |
| D630028G08Rik | 0.00420348 | 0.1151388 | -6.36742 | -2.670704152 |
| Dbf4 | 0.00329922 | 0.111481391 | 2.6835 | 1.424115888 |
| Dclk1 | 0.0000357 | 0.092008636 | 4.95448 | 2.308733647 |
| Ddah1 | 0.00856224 | 0.133080868 | 6.28506 | 2.651926517 |
| Ddo | 0.000062 | 0.09765 | -3.8995 | -1.963289904 |
| Ddx26b | 0.00101074 | 0.101611628 | -2.07477 | -1.052951819 |
| Des | 0.00292112 | 0.110861783 | -5.69198 | -2.508928864 |
| Dfna5 | 0.00100737 | 0.101633237 | 2.43649 | 1.284804301 |
| Diap3 | 0.00105393 | 0.101284459 | 2.83486 | 1.503277489 |
| Dlgap5 | 0.000248585 | 0.113667496 | 2.56175 | 1.357129691 |
| Dnajb5 | 0.00408388 | 0.114518297 | -2.31386 | -1.210299123 |
| Dnm3os | 0.00255454 | 0.11158892 | 3.77214 | 1.915383221 |
| Dpep1 | 0.00739902 | 0.129803352 | -2.75836 | -1.463807812 |
| Dpy19l1 | 0.000806469 | 0.097706821 | 2.11506 | 1.08069859 |
| Dscc1 | 0.00435719 | 0.115337382 | 2.99559 | 1.582840179 |
| Dstn | 0.00478207 | 0.116872142 | -2.55908 | -1.355626841 |
| Dtna | 0.00592452 | 0.124507148 | -5.57528 | -2.479045781 |
| E2f8 | 0.00303968 | 0.111915491 | 2.34389 | 1.228904865 |
| Ecm2 | 0.000512599 | 0.108449117 | -2.16237 | -1.112611987 |
| Ect2 | 0.00349731 | 0.11292567 | 2.16228 | 1.112553354 |
| Eef2k | 0.005809 | 0.123916591 | -2.01842 | -1.013229392 |

| | | | | |
|---|---|---|---|---|
| Eef2k | 0.000450143 | 0.110970035 | -2.71582 | -1.441387532 |
| Efhd1 | 0.0118271 | 0.144774734 | -7.96487 | -2.993654575 |
| Eltd1 | 0.0000902 | 0.1022868 | 3.4563 | 1.789228446 |
| Emb | 0.000487772 | 0.108884537 | 13.2479 | 3.727691783 |
| Emp1 | 0.00383992 | 0.114350559 | 2.1301 | 1.090921161 |
| Emr1 | 0.00857437 | 0.133050569 | 5.30869 | 2.408355898 |
| Enah | 0.00818734 | 0.131806411 | -3.41623 | -1.772406774 |
| Enpp1 | 0.00272879 | 0.11083266 | 2.56068 | 1.356526975 |
| Enpp5 | 0.00664879 | 0.126846027 | -3.84897 | -1.944471134 |
| Entpd1 | 0.00480107 | 0.117033822 | -2.25134 | -1.170780412 |
| Epha7 | 0.000107948 | 0.105528476 | -3.16795 | -1.663547498 |
| Ercc6l | 0.00909688 | 0.134391114 | 2.09534 | 1.067184362 |
| Errfi1 | 0.00432761 | 0.115634066 | -2.2051 | -1.14084145 |
| Esyt2 | 0.00617083 | 0.125677464 | -2.04303 | -1.030707656 |
| Ezh2 | 0.00579051 | 0.123895063 | 2.38523 | 1.254128387 |
| F630043A04Rik | 0.00787875 | 0.131004435 | 2.3417 | 1.227556261 |
| F730047E07Rik | 0.000749982 | 0.096208098 | 2.08669 | 1.061216289 |
| Fam105a | 0.0121424 | 0.145801372 | 5.73559 | 2.519941899 |
| Fam107a | 0.00215716 | 0.107858 | -2.48443 | -1.312914222 |
| Fam111a | 0.00937424 | 0.135453468 | 3.15649 | 1.658321181 |
| Fam129a | 0.0130049 | 0.148485266 | -2.2756 | -1.186248766 |
| Fam135a | 0.0016459 | 0.107267276 | -2.1031 | -1.072519163 |
| Fam13a | 0.00328169 | 0.111687769 | -2.95195 | -1.561668819 |
| Fam13c | 0.00389764 | 0.113798243 | -2.24718 | -1.168116275 |
| Fam150b | 0.000473403 | 0.110917149 | -2.3743 | -1.24750144 |
| Fam164a | 0.00103805 | 0.101478336 | -2.62124 | -1.390252601 |
| Fam196b | 0.0106071 | 0.139218188 | 2.1171 | 1.082089416 |
| Fam38b | 0.00584206 | 0.123783558 | 3.4616 | 1.791439026 |
| Fam60a | 0.00457671 | 0.115847972 | 2.64915 | 1.405529534 |
| Fam60a | 0.00488782 | 0.117831375 | 2.5521 | 1.35168486 |
| Fap | 0.00619847 | 0.125698587 | 3.75479 | 1.908732222 |
| Fas | 0.00967228 | 0.135814333 | -2.1487 | -1.103463084 |
| Fblim1 | 0.0123931 | 0.146271601 | -2.44271 | -1.288480425 |
| Fbn1 | 0.000944135 | 0.101004631 | 2.73229 | 1.450110617 |
| Fbxo32 | 0.00256588 | 0.111568555 | -2.22448 | -1.153465766 |
| Fcer1g | 0.0119158 | 0.145108647 | 5.385 | 2.428946345 |
| Fgf13 | 0.00869526 | 0.133248984 | -2.47526 | -1.307579944 |
| Fgf18 | 0.00872544 | 0.13342299 | 2.05494 | 1.039096271 |
| Fgfr2 | 0.000610424 | 0.098888688 | -2.66824 | -1.415887978 |
| Fibin | 0.00205508 | 0.107098379 | -2.2129 | -1.145940501 |
| Fignl1 | 0.00905274 | 0.134863468 | 2.73676 | 1.452468923 |
| Flna | 0.0112508 | 0.142583898 | -3.37417 | -1.754533544 |
| Fmo1 | 0.0044243 | 0.115709322 | -3.096 | -1.630402863 |
| Fndc1 | 0.00934605 | 0.135183938 | 2.6672 | 1.415326009 |
| Foxc1 | 0.00649552 | 0.126562194 | -2.70108 | -1.433537467 |
| Foxd1 | 0.0000129 | 0.045714375 | -2.50627 | -1.325539348 |
| Fry | 0.00272106 | 0.11083628 | -3.7617 | -1.911386178 |
| Fscn1 | 0.00789968 | 0.130968379 | 3.13375 | 1.647890091 |
| Fstl1 | 0.0051923 | 0.120954565 | 2.15649 | 1.108685026 |
| Ftl1 | 0.00205011 | 0.107233613 | 3.03917 | 1.603677376 |
| Ftl1 | 0.00274271 | 0.110763288 | 2.46464 | 1.301376933 |
| Ftl1 | 0.00189738 | 0.106939807 | 2.34166 | 1.227531617 |

| | | | | |
|---|---|---|---|---|
| Ftl2 | 0.00260117 | 0.111394516 | 2.40333 | 1.265034758 |
| Ftl2 | 0.00228549 | 0.109080205 | 2.28461 | 1.191947907 |
| Fxyd1 | 0.00464219 | 0.11605475 | -2.1406 | -1.098014433 |
| Fxyd5 | 0.00498306 | 0.118714076 | 2.56686 | 1.360004612 |
| Fyco1 | 0.000697945 | 0.095128561 | -2.34453 | -1.22929779 |
| Fzd7 | 0.00216191 | 0.107905191 | -2.2025 | -1.139140468 |
| Galc | 0.000128492 | 0.110386309 | 2.35448 | 1.235408468 |
| Gba | 0.00966973 | 0.135913161 | 2.07476 | 1.052944461 |
| Gda | 0.0120472 | 0.145645254 | 2.89409 | 1.533109787 |
| Gen1 | 0.00363369 | 0.112216897 | 2.31018 | 1.208005265 |
| Gjc1 | 0.00463788 | 0.116049336 | -2.43244 | -1.28240363 |
| Gkap1 | 0.00733113 | 0.130061036 | -2.02437 | -1.017475463 |
| Gkn3 | 0.0105114 | 0.138926895 | -7.7212 | -2.948820038 |
| Glb1l2 | 0.00783583 | 0.130904997 | -3.67481 | -1.877669196 |
| Gm10118 | 0.00634003 | 0.125956447 | -4.33446 | -2.11585381 |
| Gm106 | 0.00563996 | 0.122994512 | -3.56647 | -1.834498343 |
| Gm10743 | 0.00773034 | 0.13060497 | -2.3369 | -1.224593729 |
| Gm10850 | 0.0100057 | 0.137299901 | -3.6133 | -1.853318713 |
| Gm11428 | 0.0115796 | 0.143856994 | 7.37795 | 2.883220012 |
| Gm11711 | 0.00477777 | 0.116968722 | 2.185 | 1.12763328 |
| Gm11711 | 0.0067323 | 0.127666023 | 2.16961 | 1.117435733 |
| Gm447 | 0.0116972 | 0.144118044 | -2.12366 | -1.086550265 |
| Gm5820 | 0.000878799 | 0.098474117 | -2.78059 | -1.475394663 |
| Gm885 | 0.0111953 | 0.142134687 | 4.68634 | 2.228461627 |
| Gnai1 | 0.0112578 | 0.142608861 | -3.91921 | -1.97056549 |
| Gp49a | 0.0100925 | 0.137625 | 8.37965 | 3.066889987 |
| Gpm6b | 0.00127209 | 0.10669749 | 2.32804 | 1.219115847 |
| Gpr137b-ps | 0.00329747 | 0.1116885 | 2.64119 | 1.401188089 |
| Gpr183 | 0.00404752 | 0.114518156 | 2.25246 | 1.171501486 |
| Gpr21 | 0.0017941 | 0.107988822 | -3.08934 | -1.627301928 |
| Gpx3 | 0.00663933 | 0.126921784 | 4.27875 | 2.097189387 |
| Greb1l | 0.000542155 | 0.102467295 | -2.62783 | -1.393872404 |
| Grip2 | 0.000174157 | 0.123433774 | -3.05909 | -1.613105198 |
| Gsg2 | 0.00480055 | 0.117121852 | 2.56334 | 1.35802485 |
| Gsn | 0.00663975 | 0.126844281 | -2.54358 | -1.346859249 |
| Gsta4 | 0.000647078 | 0.09456011 | -3.24243 | -1.697073873 |
| Gstk1 | 0.0082685 | 0.132361364 | -2.12193 | -1.085377311 |
| Gstm1 | 0.00288386 | 0.110632518 | -2.67437 | -1.419198456 |
| Gstm3 | 0.00645679 | 0.126415743 | -2.01277 | -1.009178708 |
| Gsto1 | 0.00476406 | 0.117138856 | 2.05014 | 1.035722432 |
| Gusb | 0.00230218 | 0.109141811 | 2.07551 | 1.053465883 |
| H19 | 0.00654159 | 0.126589813 | 8.12543 | 3.022444163 |
| H2-Aa | 0.00697743 | 0.129034664 | 5.57971 | 2.480190141 |
| H2-Ab1 | 0.0113762 | 0.142705872 | 5.04491 | 2.334828532 |
| H2afz | 0.000516013 | 0.107565945 | 2.17565 | 1.121446487 |
| H2afz | 0.000590769 | 0.100289228 | 2.09244 | 1.065186255 |
| H2-DMa | 0.00829446 | 0.132477713 | 5.1875 | 2.375039431 |
| H2-Eb1 | 0.0119786 | 0.145373848 | 5.73792 | 2.520527854 |
| Havcr2 | 0.00978067 | 0.136592116 | 4.65458 | 2.218650994 |
| Hcls1 | 0.00943804 | 0.135409127 | 3.29022 | 1.718184053 |
| Hells | 0.00000379 | 0.026861625 | 6.17891 | 2.62735236 |
| Hhip | 0.000427838 | 0.109272138 | -5.23825 | -2.389080763 |

| | | | | |
|---|---|---|---|---|
| Higd1a | 0.000451467 | 0.110336978 | -2.70321 | -1.434671892 |
| Higd1a | 0.00318093 | 0.11230307 | -3.11785 | -1.640550797 |
| Higd1b | 0.00682096 | 0.128062395 | -3.05958 | -1.61333471 |
| Hist1h2bk | 0.00290699 | 0.110770385 | 2.14659 | 1.102046662 |
| Hist1h2bl | 0.00381289 | 0.113904564 | 2.22036 | 1.150793608 |
| Hist1h2bn | 0.0025511 | 0.111956169 | 2.39506 | 1.260061798 |
| Hist2h2be | 0.00268283 | 0.11071067 | -3.05388 | -1.610640246 |
| Hlf | 0.00000826 | 0.0468342 | -3.27716 | -1.712447328 |
| Hmgb2 | 0.0022018 | 0.108937225 | 2.69974 | 1.432820474 |
| Hmgb2 | 0.0030132 | 0.111811806 | 2.66824 | 1.415888438 |
| Hmgb2 | 0.0028673 | 0.110746533 | 2.59583 | 1.376195905 |
| Hmgb2 | 0.00246973 | 0.111847996 | 2.49965 | 1.321726103 |
| Hmmr | 0.00148927 | 0.106618193 | 3.84632 | 1.943478795 |
| Hn1 | 0.000392452 | 0.110158556 | 2.42624 | 1.278722267 |
| Hn1l | 0.00745616 | 0.129523368 | 2.54699 | 1.348793295 |
| Hpse2 | 0.00977616 | 0.13659642 | -7.5001 | -2.906905023 |
| Hrc | 0.000247495 | 0.116941388 | -5.87318 | -2.554137719 |
| Hspa1a | 0.00607559 | 0.124994903 | -2.3684 | -1.243909258 |
| Hspa1a | 0.00171465 | 0.107544972 | -3.42669 | -1.776814726 |
| Hspa1a | 0.00144691 | 0.106822652 | -4.16386 | -2.057920198 |
| Hspa1l | 0.0105561 | 0.138934742 | -2.52171 | -1.334399845 |
| Hspa2 | 0.00583898 | 0.123810832 | -2.05534 | -1.039378572 |
| Hspb1 | 0.00539904 | 0.121962378 | -4.01465 | -2.005272575 |
| Hspb1 | 0.00588476 | 0.124316651 | -4.23328 | -2.081778654 |
| Hspb6 | 0.00456897 | 0.115962667 | -4.84326 | -2.275981946 |
| Hspb7 | 0.00937998 | 0.135329482 | -2.82896 | -1.500270936 |
| Hsph1 | 0.00449456 | 0.115942471 | -2.47438 | -1.307069375 |
| Htr2a | 0.00348373 | 0.112872852 | -2.62708 | -1.393459227 |
| Hyal1 | 0.00622281 | 0.125742454 | -2.53849 | -1.343974155 |
| Id2 | 0.0111852 | 0.142133761 | -2.64481 | -1.403164061 |
| Igf1 | 0.00561843 | 0.122903156 | 3.00016 | 1.585039442 |
| Il10rb | 0.000822996 | 0.097623166 | 3.22763 | 1.690475204 |
| Il17ra | 0.010178 | 0.137928442 | 2.30622 | 1.205530144 |
| Il2rg | 0.00615616 | 0.125649486 | 9.67546 | 3.274330253 |
| Il33 | 0.00267288 | 0.110622114 | 4.40155 | 2.138011656 |
| Il4ra | 0.000643939 | 0.095579427 | 3.31451 | 1.728795606 |
| Inmt | 0.00145194 | 0.106638598 | -17.3064 | -4.113231056 |
| Inpp4a | 0.00253909 | 0.111948991 | -3.9413 | -1.978673789 |
| Inpp5a | 0.0129462 | 0.148053558 | -2.63506 | -1.397835812 |
| Irf8 | 0.0109234 | 0.141019303 | 2.18354 | 1.12666896 |
| Itga1 | 0.00689735 | 0.128560074 | -2.33356 | -1.222531889 |
| Itga11 | 0.00234541 | 0.10990475 | 3.06993 | 1.61820576 |
| Itga6 | 0.00579072 | 0.123806118 | 2.81783 | 1.494584576 |
| Itga9 | 0.00132013 | 0.104833853 | -3.46501 | -1.792856795 |
| Itgam | 0.003995 | 0.113941901 | 2.77927 | 1.474705996 |
| Itgax | 0.0097241 | 0.1361374 | 4.75263 | 2.24872609 |
| Itgb2 | 0.0081331 | 0.13145575 | 3.20819 | 1.681759586 |
| Itm2a | 0.0074373 | 0.129513179 | 3.22311 | 1.688453426 |
| Itpk1 | 0.00531893 | 0.121312683 | -2.52196 | -1.334545375 |
| Itpr1 | 0.00168925 | 0.107377214 | -2.35385 | -1.235022072 |
| Itpr1 | 0.00109502 | 0.101450382 | -2.61835 | -1.388657623 |
| Ivns1abp | 0.0130991 | 0.149140355 | 2.06647 | 1.04716842 |

| | | | | |
|---|---|---|---|---|
| Jph1 | 0.0117733 | 0.144553077 | -2.86032 | -1.51617339 |
| Jph2 | 0.0025594 | 0.111629215 | -3.65999 | -1.871838599 |
| Kank1 | 0.00818279 | 0.131957962 | -3.02856 | -1.598631667 |
| Kcna5 | 0.00100692 | 0.10195065 | -3.86316 | -1.94978391 |
| Kcnmb1 | 0.00602494 | 0.124858954 | -4.40037 | -2.137622407 |
| Kif11 | 0.00164237 | 0.107780531 | 6.76007 | 2.757038186 |
| Kif15 | 0.00387384 | 0.114042953 | 2.61899 | 1.389010551 |
| Kif1c | 0.00193799 | 0.107729444 | -2.08045 | -1.056896339 |
| Kif20b | 0.00310945 | 0.112583534 | 3.32425 | 1.733028884 |
| Kif23 | 0.000927244 | 0.100333463 | 3.1886 | 1.672923127 |
| Kif4 | 0.00341957 | 0.112074924 | 3.4671 | 1.793729448 |
| Klk10 | 0.000911476 | 0.100156374 | -2.74634 | -1.457510147 |
| Klrb1b | 0.00544355 | 0.122382746 | 4.81855 | 2.268599075 |
| Kng2 | 0.0101292 | 0.137661946 | 2.72092 | 1.444094539 |
| Kntc1 | 0.00168838 | 0.107563085 | 2.04262 | 1.030420836 |
| Kynu | 0.00502281 | 0.119160388 | 4.03964 | 2.01422673 |
| Lamb1 | 0.000767999 | 0.095077605 | 2.8947 | 1.533413838 |
| Lancl3 | 0.00110774 | 0.101632456 | -2.18772 | -1.129427745 |
| Laptm5 | 0.00261782 | 0.110768951 | 4.72667 | 2.240824144 |
| Lbp | 0.00384432 | 0.113883461 | 2.06366 | 1.045205298 |
| Lcp1 | 0.00518909 | 0.121178502 | 7.27001 | 2.861957349 |
| Lcp2 | 0.00437029 | 0.11546852 | 4.18582 | 2.065510274 |
| Ldb3 | 0.00415286 | 0.114750079 | -3.1202 | -1.64163975 |
| Ldhb | 0.00326735 | 0.112142097 | -3.84893 | -1.944460028 |
| Lepr | 0.00777084 | 0.130821445 | -2.65624 | -1.409381694 |
| Leprel1 | 0.0122156 | 0.145754318 | -3.88406 | -1.957562977 |
| Lgals3 | 0.00182946 | 0.1067185 | 2.5793 | 1.366979584 |
| Lgmn | 0.000473443 | 0.110017287 | 3.62193 | 1.856758664 |
| Lgr4 | 0.00331726 | 0.111295054 | -2.19163 | -1.132002353 |
| Lilrb4 | 0.0106142 | 0.139118155 | 6.97063 | 2.801289052 |
| Limch1 | 0.00361654 | 0.112053452 | -2.0811 | -1.057346627 |
| Lims2 | 0.0105423 | 0.138946632 | -2.83448 | -1.503081623 |
| Lipa | 0.000544967 | 0.102316652 | 2.92305 | 1.547474507 |
| Lmnb1 | 0.00214229 | 0.107875527 | 3.02493 | 1.596901757 |
| Lmod1 | 0.00897018 | 0.134695235 | -5.83161 | -2.543896186 |
| Lpar6 | 0.00335787 | 0.111470275 | 2.15899 | 1.110356561 |
| Lpp | 0.00899343 | 0.134758848 | -2.52563 | -1.336642628 |
| Lpp | 0.000957326 | 0.10051923 | -2.70628 | -1.436310782 |
| Lrrc55 | 0.0130646 | 0.148986891 | 2.20212 | 1.138893088 |
| Lrrn1 | 0.000393441 | 0.109353454 | -4.46678 | -2.159234663 |
| Lrrtm3 | 0.0131728 | 0.149439328 | -4.75052 | -2.248087301 |
| Lsp1 | 0.000705013 | 0.094278861 | 2.9396 | 1.555619857 |
| Ly9 | 0.00319178 | 0.112406165 | 3.44981 | 1.786516907 |
| Lyve1 | 0.0102838 | 0.138304426 | -2.78397 | -1.477140745 |
| Mad2l1 | 0.000376151 | 0.107715968 | 2.54888 | 1.349863454 |
| Mafb | 0.0122649 | 0.146035244 | 2.17068 | 1.118147061 |
| Marcks | 0.00411511 | 0.114600558 | 2.1576 | 1.109427427 |
| Marveld1 | 0.00383436 | 0.114305054 | -2.98214 | -1.576346537 |
| Mastl | 0.0110505 | 0.141245119 | 3.14864 | 1.654728816 |
| Me1 | 0.00693307 | 0.128633858 | -2.09469 | -1.066738594 |
| Me1 | 0.0078689 | 0.130994313 | -2.47191 | -1.305627904 |
| Mefv | 0.0107279 | 0.140025767 | 2.47355 | 1.306583062 |

| | | | | |
|---|---|---|---|---|
| Mertk | 0.00320054 | 0.112296175 | -2.42468 | -1.277792809 |
| Mfap5 | 0.000100359 | 0.10537695 | 2.57022 | 1.361891853 |
| Mfhas1 | 0.00256888 | 0.111357413 | -2.01724 | -1.012379347 |
| Mfsd1 | 0.000209088 | 0.118552896 | 2.37669 | 1.24895374 |
| Mgll | 0.00838262 | 0.132689714 | -2.33449 | -1.223104327 |
| Mir143 | 0.00804102 | 0.130862754 | -5.78476 | -2.532257262 |
| Mir145 | 0.00011792 | 0.107839742 | -6.42582 | -2.683882069 |
| Mir29b-2 | 0.00554935 | 0.123101778 | -4.34516 | -2.11941007 |
| Mir29c | 0.00321503 | 0.112248892 | -7.53471 | -2.913553174 |
| Mki67 | 0.00397461 | 0.114280115 | 4.71531 | 2.237352622 |
| Mllt3 | 0.000812758 | 0.097634277 | -2.34482 | -1.22947707 |
| Mmp12 | 0.00291489 | 0.110922324 | 15.7816 | 3.980171574 |
| Mmp14 | 0.00180854 | 0.107941282 | 4.77538 | 2.25561554 |
| Mmp2 | 0.0109687 | 0.141090129 | 3.29583 | 1.72064183 |
| Mmp3 | 0.000270075 | 0.11779425 | 6.27574 | 2.649785584 |
| Mpeg1 | 0.0054105 | 0.121929789 | 8.42582 | 3.074817096 |
| Mpp6 | 0.000193628 | 0.12198564 | 2.78319 | 1.4767394 |
| Mpp7 | 0.00261344 | 0.110748915 | -3.87189 | -1.95303685 |
| Mrc2 | 0.000337287 | 0.107439174 | 2.56007 | 1.356183258 |
| Mrgprh | 0.00260876 | 0.11155105 | -6.64565 | -2.732413866 |
| Mrpl48 | 0.00143763 | 0.106973256 | -2.23068 | -1.157482905 |
| Mrps6 | 0.00165444 | 0.107330375 | -2.17691 | -1.122280875 |
| Mrvi1 | 0.0109453 | 0.141109256 | -2.37745 | -1.249414075 |
| Ms4a7 | 0.00353643 | 0.113157777 | 13.0494 | 3.705911569 |
| Msr1 | 0.0108303 | 0.140779003 | 6.60798 | 2.72420932 |
| Mtss1l | 0.00212218 | 0.108014009 | -3.94012 | -1.97824171 |
| Mtus2 | 0.0025623 | 0.111584032 | -2.17052 | -1.118044131 |
| Mum1l1 | 0.00148655 | 0.106692892 | -2.80544 | -1.488228374 |
| Myl4 | 0.000717102 | 0.09499926 | -4.10447 | -2.03719485 |
| Mylk | 0.00761681 | 0.13063313 | -3.19295 | -1.674889946 |
| Myo10 | 0.00323635 | 0.112301741 | 3.03363 | 1.601045136 |
| Myo5a | 0.00337367 | 0.111733113 | 2.35761 | 1.237325085 |
| Nalcn | 0.00570561 | 0.123288143 | -2.77271 | -1.471292669 |
| Nap1l5 | 0.000644069 | 0.095100813 | -2.19167 | -1.132033972 |
| Nav3 | 0.00342912 | 0.112258143 | 2.16783 | 1.116251626 |
| Ncapg | 0.00219742 | 0.109101326 | 4.88734 | 2.289049473 |
| Ncapg2 | 0.00266761 | 0.110727296 | 2.81735 | 1.494338801 |
| Ncaph | 0.00863356 | 0.133167261 | 2.49011 | 1.316209474 |
| Nckap1l | 0.00471938 | 0.116647274 | 5.46073 | 2.449093826 |
| Ndc80 | 0.000213831 | 0.116579016 | 3.94437 | 1.979794889 |
| Neurl3 | 0.0125055 | 0.146500382 | 2.0326 | 1.023326332 |
| Nexn | 0.0103528 | 0.13837901 | -3.25958 | -1.704686042 |
| Nfia | 0.0039144 | 0.114052662 | -2.02728 | -1.019541773 |
| Nfia | 0.00948192 | 0.135490137 | -2.03878 | -1.027704371 |
| Nid2 | 0.0094969 | 0.135431144 | 2.17198 | 1.119010819 |
| Nkd1 | 0.00274721 | 0.110472913 | -4.18313 | -2.064579479 |
| Nov | 0.000339909 | 0.107071335 | -2.24743 | -1.168275141 |
| Npc2 | 0.00244843 | 0.111776152 | 2.13269 | 1.092674276 |
| Npnt | 0.00358488 | 0.112299832 | -4.29882 | -2.10394055 |
| Nr3c2 | 0.0038866 | 0.113827593 | -3.0716 | -1.618991347 |
| Nrip2 | 0.00361602 | 0.112282768 | -2.4546 | -1.295489197 |
| Nrp2 | 0.000436932 | 0.110598413 | 2.32389 | 1.216541781 |

| | | | | |
|---|---|---|---|---|
| Nt5dc2 | 0.00460652 | 0.115878298 | 2.6667 | 1.415055533 |
| Nuf2 | 0.00116363 | 0.102769192 | 4.85802 | 2.280368429 |
| Nusap1 | 0.00652813 | 0.126588567 | 2.10424 | 1.073299261 |
| Ogn | 0.0000361 | 0.08528625 | -3.53196 | -1.820468567 |
| Optc | 0.00881024 | 0.133996944 | -2.13759 | -1.095983806 |
| Ostb | 0.00559704 | 0.123004716 | -2.11708 | -1.082074891 |
| P2rx4 | 0.0056536 | 0.123102581 | 2.3803 | 1.251143414 |
| P4ha3 | 0.00153679 | 0.107310336 | 6.91871 | 2.790503071 |
| Pak1 | 0.00421424 | 0.115210901 | 3.40738 | 1.76866285 |
| Palld | 0.00391577 | 0.113975441 | -2.79256 | -1.48158572 |
| Pappa2 | 0.00982655 | 0.136694157 | -2.13259 | -1.092610895 |
| Parm1 | 0.00357889 | 0.112735035 | -3.83519 | -1.939299573 |
| Pbk | 0.00289541 | 0.110626514 | 3.69722 | 1.886440892 |
| Pcolce2 | 0.00266489 | 0.110776586 | -4.71868 | -2.238381115 |
| Pcp4l1 | 0.00899361 | 0.134690356 | -10.5799 | -3.403260982 |
| Pdcl3 | 0.00153355 | 0.108149608 | -2.31667 | -1.212052737 |
| Pde4dip | 0.00307936 | 0.111922892 | -2.63542 | -1.398033509 |
| Pde5a | 0.0114282 | 0.142789542 | -3.82453 | -1.935282668 |
| Pdgfd | 0.00413645 | 0.114856374 | -2.68667 | -1.425815155 |
| Pdlim3 | 0.0120409 | 0.145631192 | -8.82932 | -3.142302398 |
| Pdpn | 0.00638826 | 0.126471488 | 3.00218 | 1.586010478 |
| Pgam2 | 0.00001 | 0.04725 | -3.6149 | -1.853954827 |
| Pgm5 | 0.0015416 | 0.106596 | -5.699 | -2.510711922 |
| Phex | 0.00196771 | 0.107484737 | -4.77318 | -2.254950306 |
| Phtf2 | 0.00841913 | 0.132748796 | -2.53863 | -1.344051065 |
| Pik3cg | 0.0050441 | 0.119465526 | 2.65652 | 1.409537574 |
| Pip5k1b | 0.00438857 | 0.115306728 | -3.8197 | -1.933457487 |
| Pira1 | 0.0103191 | 0.138319851 | 3.90338 | 1.964723918 |
| Pkhd1l1 | 0.00140299 | 0.106921415 | -6.4675 | -2.693210482 |
| Pkia | 0.00807961 | 0.13111445 | -3.30438 | -1.724382619 |
| Pkp4 | 0.000626723 | 0.09816352 | -3.64923 | -1.867594251 |
| Pla2g7 | 0.00348918 | 0.11292038 | 3.45515 | 1.788748344 |
| Plac8 | 0.00973557 | 0.136163498 | 8.51304 | 3.089674409 |
| Plau | 0.00174056 | 0.107505176 | 2.46173 | 1.299672537 |
| Plaur | 0.00839123 | 0.132677842 | 2.04491 | 1.032037349 |
| Plbd1 | 0.0109549 | 0.141168825 | 6.13446 | 2.616936353 |
| Plcb4 | 0.00438119 | 0.11554115 | -3.27685 | -1.712310224 |
| Plin2 | 0.0104381 | 0.138734241 | 2.56299 | 1.35782785 |
| Plin4 | 0.00828474 | 0.132397057 | -3.04495 | -1.606421251 |
| Plk1 | 0.00259414 | 0.111430105 | 2.06404 | 1.04547093 |
| Plk2 | 0.0114947 | 0.143304637 | 2.48913 | 1.31564158 |
| Pln | 0.00631687 | 0.125760719 | -6.53072 | -2.707237094 |
| Pls3 | 0.00697149 | 0.129093234 | -2.12531 | -1.087672047 |
| Plscr1 | 0.00358985 | 0.111837635 | 2.3211 | 1.21480868 |
| Plscr1 | 0.0078526 | 0.130953653 | 2.26886 | 1.18196759 |
| Pnck | 0.00576971 | 0.123823829 | -2.50414 | -1.32431412 |
| Pnldc1 | 0.00426774 | 0.115780315 | -2.03454 | -1.02469852 |
| Pnpla2 | 0.00241659 | 0.111218063 | -2.36233 | -1.240210116 |
| Podn | 0.000143578 | 0.110011792 | -2.03548 | -1.025367904 |
| Postn | 0.000541457 | 0.103022188 | 6.9591 | 2.798900739 |
| Ppargc1a | 0.00725309 | 0.129568432 | -7.77333 | -2.958532709 |
| Ppic | 0.000485326 | 0.110071937 | 2.02449 | 1.017558517 |

| | | | | |
|---|---|---|---|---|
| Ppm1e | 0.000634044 | 0.096123783 | -2.41591 | -1.272565679 |
| Ppp1cb | 0.00640093 | 0.126369335 | -2.08251 | -1.058322735 |
| Ppp1r12a | 0.00547166 | 0.12262574 | -2.68159 | -1.423089004 |
| Ppp1r12b | 0.00332519 | 0.111429239 | -3.89003 | -1.959778061 |
| Ppp1r9a | 0.00354256 | 0.112971402 | -2.30727 | -1.206188997 |
| Prc1 | 0.00111899 | 0.101677457 | 4.79134 | 2.260429193 |
| Prcp | 0.0027565 | 0.109911076 | 2.87796 | 1.525046541 |
| Prdm1 | 0.0100792 | 0.137509779 | 2.87794 | 1.525036515 |
| Prdm16 | 0.00769272 | 0.130513831 | -2.35402 | -1.235127348 |
| Prdm6 | 0.00512579 | 0.120494317 | -2.21837 | -1.149501386 |
| Prg4 | 0.00518961 | 0.1210909 | 5.8159 | 2.540002463 |
| Prkag2 | 0.00221904 | 0.108652477 | -2.44915 | -1.29228086 |
| Prkg1 | 0.00398786 | 0.114197809 | -3.53762 | -1.822778695 |
| Prnd | 0.00612947 | 0.125647487 | 12.8321 | 3.681685385 |
| Prr11 | 0.0118288 | 0.144733051 | 3.57845 | 1.839334822 |
| Prss23 | 0.0031466 | 0.112350264 | -2.00409 | -1.002946104 |
| Ptger3 | 0.000419271 | 0.109048925 | -6.89735 | -2.786044348 |
| Ptgis | 0.011345 | 0.142693323 | -3.09911 | -1.631855231 |
| Ptpla | 0.0105894 | 0.139114685 | -2.13419 | -1.093691216 |
| Ptpla | 0.00740632 | 0.129770811 | -2.23679 | -1.161430583 |
| Ptprb | 0.00535753 | 0.121411651 | -2.06295 | -1.04470803 |
| Ptprc | 0.00768906 | 0.130529851 | 6.73352 | 2.751360882 |
| Ptpre | 0.00586081 | 0.123995495 | 2.42058 | 1.275352776 |
| Ptprz1 | 0.000975608 | 0.101313138 | -12.6881 | -3.665400605 |
| Pvt1 | 0.00051986 | 0.106797326 | 2.08744 | 1.06173473 |
| Pycard | 0.0070527 | 0.129329913 | 2.3657 | 1.242267134 |
| Pygb | 0.00804457 | 0.130845416 | -2.47499 | -1.307419256 |
| Pygm | 0.0054185 | 0.122013086 | -2.84785 | -1.509873528 |
| Rab31 | 0.00368884 | 0.112450123 | 2.25737 | 1.174642907 |
| Rad51 | 0.00688578 | 0.128513406 | 2.99991 | 1.584919219 |
| Raet1d | 0.00434555 | 0.115677317 | 2.24176 | 1.164631833 |
| Ralgapa2 | 0.00110626 | 0.101826205 | -2.38565 | -1.254378861 |
| Ramp1 | 0.000805854 | 0.098051334 | -2.09357 | -1.065968189 |
| Rasa3 | 0.000859807 | 0.097113659 | 2.19993 | 1.137457619 |
| Rasgrf2 | 0.00463513 | 0.116082982 | -2.4182 | -1.2739361 |
| Rasgrp2 | 0.000682833 | 0.094430808 | -2.28017 | -1.189143993 |
| Rasl12 | 0.0046527 | 0.115908651 | -3.41016 | -1.769836345 |
| Rassf3 | 0.0076293 | 0.130610299 | -2.25279 | -1.171716137 |
| Rbp4 | 0.00529667 | 0.121194992 | -5.99861 | -2.584630719 |
| Rbpms | 0.00646149 | 0.126420457 | -3.41335 | -1.771189927 |
| Rbpms2 | 0.0071644 | 0.129205305 | -3.41164 | -1.770466218 |
| Reck | 0.00725374 | 0.129498444 | -2.00854 | -1.00614452 |
| Retnla | 0.00925534 | 0.135182323 | -13.0958 | -3.71103153 |
| Rgs4 | 0.00740184 | 0.129772519 | -8.24209 | -3.043015562 |
| Rgs5 | 0.00526671 | 0.12089978 | -2.25705 | -1.174435779 |
| Rgs5 | 0.00708624 | 0.12952605 | -3.1222 | -1.642562853 |
| Rgs7bp | 0.00466953 | 0.116123838 | -4.82008 | -2.269053171 |
| Rnd3 | 0.00312726 | 0.112367327 | -2.41191 | -1.270176661 |
| Rnf152 | 0.00126683 | 0.106888781 | -2.26354 | -1.178583659 |
| Rnf213 | 0.0124892 | 0.146551664 | 3.963 | 1.986592967 |
| Rock1 | 0.000499767 | 0.10898765 | -2.29308 | -1.197288954 |
| Rplp1 | 0.00800774 | 0.131073573 | 2.11419 | 1.080105036 |

| | | | | |
|---|---|---|---|---|
| Rras | 0.00104486 | 0.101098229 | -2.54529 | -1.347832023 |
| Rrm1 | 0.000948308 | 0.101069668 | 2.00221 | 1.001593298 |
| Rrm2 | 0.000739692 | 0.096193891 | 2.49134 | 1.316921924 |
| Runx1 | 0.000524895 | 0.105537399 | 6.67875 | 2.739578112 |
| Ryr3 | 0.00255235 | 0.111837902 | -2.36355 | -1.240956688 |
| S1pr3 | 0.00029278 | 0.11067084 | -6.3688 | -2.671025705 |
| Saa3 | 0.00820786 | 0.131912036 | 26.1968 | 4.711318689 |
| Samd12 | 0.00931997 | 0.135289887 | -2.04589 | -1.032728083 |
| Samd4 | 0.00160323 | 0.107705143 | -2.6062 | -1.38194933 |
| Scd1 | 0.00962783 | 0.135930767 | -4.48072 | -2.163733276 |
| Scrn3 | 0.0104784 | 0.138749482 | -2.09878 | -1.06954877 |
| Selplg | 0.00597952 | 0.124922175 | 2.74818 | 1.458476501 |
| Sema3c | 0.00264338 | 0.111186681 | -4.39419 | -2.135598693 |
| Sema5a | 0.0123396 | 0.146371406 | -2.40397 | -1.265417397 |
| Serpina3n | 0.000234895 | 0.114815056 | 5.87639 | 2.554930147 |
| Serpini1 | 0.0121138 | 0.145673905 | -2.87154 | -1.521821313 |
| Sfpi1 | 0.00677341 | 0.127846986 | 3.21117 | 1.683099044 |
| Sfrp1 | 0.00228485 | 0.109233554 | 4.26426 | 2.092295405 |
| Sgol2 | 0.00565258 | 0.123174975 | 3.16747 | 1.663330955 |
| Sh2d1b1 | 0.00951241 | 0.135447927 | 10.0319 | 3.326522967 |
| Sh3bgr | 0.00899237 | 0.13481422 | -5.39695 | -2.432143073 |
| Sh3bgrl3 | 0.00942638 | 0.135378862 | 2.66758 | 1.415531538 |
| Sh3bp2 | 0.0104936 | 0.138821073 | 2.47279 | 1.306139725 |
| Shcbp1 | 0.00410225 | 0.114354757 | 4.45821 | 2.156464575 |
| Sirpa | 0.00164428 | 0.107408613 | 3.14141 | 1.651412248 |
| Sirpb1a | 0.00602855 | 0.124842507 | 8.9351 | 3.159483876 |
| Sirpb1a | 0.00734905 | 0.130134646 | 7.5329 | 2.913205378 |
| Sirpb1b | 0.00641401 | 0.126363574 | 7.56502 | 2.919343897 |
| Sirpb1b | 0.00911896 | 0.134437086 | 7.54635 | 2.915779014 |
| Skap2 | 0.0108062 | 0.140659215 | 2.26902 | 1.182069326 |
| Slamf9 | 0.0114158 | 0.142886503 | 3.83999 | 1.941102554 |
| Slc16a7 | 0.00356445 | 0.112781426 | -2.87579 | -1.523956407 |
| Slc22a3 | 0.00165126 | 0.107369773 | -3.74411 | -1.90462374 |
| Slc25a23 | 0.0101732 | 0.137995321 | -2.83602 | -1.503866979 |
| Slc2a4 | 0.00338041 | 0.111825698 | -2.65119 | -1.406640492 |
| Slc38a1 | 0.0104747 | 0.138960105 | 4.28408 | 2.098985421 |
| Slc38a11 | 0.00231618 | 0.10889503 | -7.85734 | -2.974035707 |
| Slc4a3 | 0.000704943 | 0.094716275 | -8.88475 | -3.151336402 |
| Slc4a4 | 0.00101997 | 0.101460174 | -2.15768 | -1.109483266 |
| Slc5a3 | 0.0065982 | 0.126905678 | -2.75302 | -1.461016929 |
| Slc7a11 | 0.00314178 | 0.112319625 | 3.15435 | 1.657342747 |
| Slc7a7 | 0.00788118 | 0.130968026 | 2.54395 | 1.347070315 |
| Slc9a7 | 0.0100215 | 0.137250978 | 2.32681 | 1.21835341 |
| Slit3 | 0.00106457 | 0.100938326 | -2.61173 | -1.385005649 |
| Smarcd3 | 0.000618896 | 0.098571357 | -3.52773 | -1.818742213 |
| Smc2 | 0.00453548 | 0.115838611 | 2.44236 | 1.288275867 |
| Snord33 | 0.00446325 | 0.116085447 | 2.23888 | 1.162777204 |
| Snrpn | 0.00976085 | 0.136449752 | -2.30047 | -1.201927912 |
| Soat1 | 0.0059186 | 0.124659963 | 2.32302 | 1.216001575 |
| Sorbs2 | 0.0055787 | 0.122792038 | -6.7309 | -2.750804687 |
| Sorl1 | 0.0000597 | 0.105780938 | -2.21459 | -1.147035959 |
| Sort1 | 0.00470556 | 0.116610687 | -3.37168 | -1.753467868 |

| | | | | |
|---|---|---|---|---|
| Sox6 | 0.00959896 | 0.135658283 | -3.17746 | -1.667877569 |
| Spc24 | 0.00238366 | 0.110239414 | 2.24865 | 1.169059125 |
| Spc25 | 0.00662798 | 0.126961644 | 3.55787 | 1.831013797 |
| Speg | 0.0119613 | 0.145350559 | -2.31564 | -1.211411168 |
| Spon1 | 0.00168809 | 0.107786828 | 2.05024 | 1.035792801 |
| Spp1 | 0.00000226 | 0.0320355 | 32.9054 | 5.040252454 |
| Srpx2 | 0.0020351 | 0.108245938 | 3.03881 | 1.603506474 |
| St5 | 0.00431285 | 0.115457316 | -2.10708 | -1.075243353 |
| St8sia4 | 0.00420196 | 0.115208478 | 3.70291 | 1.888659485 |
| Stac | 0.00605723 | 0.125253443 | -2.4252 | -1.278104171 |
| Stbd1 | 0.0108002 | 0.140645691 | -2.29259 | -1.196981322 |
| Stk17b | 0.0116028 | 0.143892992 | 2.28803 | 1.194105968 |
| Stmn1 | 0.000854397 | 0.098065405 | 4.18763 | 2.066133978 |
| Stmn1 | 0.00092603 | 0.100586017 | 4.17133 | 2.06050745 |
| Strbp | 0.00916371 | 0.134606828 | -2.00151 | -1.00109109 |
| Sult1a1 | 0.00997052 | 0.137415772 | -2.27781 | -1.187648003 |
| Susd2 | 0.000500617 | 0.108339633 | -3.16031 | -1.660064507 |
| Sykb | 0.00533833 | 0.121462003 | 2.31927 | 1.213670782 |
| Syne2 | 0.0123758 | 0.146250075 | -2.21085 | -1.144600254 |
| Synpo2 | 0.0123359 | 0.146388767 | -9.3417 | -3.22368373 |
| Sytl5 | 0.000587747 | 0.100377274 | -4.28624 | -2.099710869 |
| Sytl5 | 0.0000576 | 0.108864 | -5.34742 | -2.418843536 |
| Tacc3 | 0.00159159 | 0.107946355 | 3.03493 | 1.601663241 |
| Tcap | 0.0104087 | 0.138538331 | -2.00521 | -1.003752999 |
| Tceal3 | 0.0074436 | 0.129543315 | -2.30267 | -1.203309225 |
| Tcf15 | 0.00158274 | 0.107862209 | -2.19407 | -1.133609468 |
| Tcirg1 | 0.0019986 | 0.107719221 | 2.0686 | 1.048654702 |
| Tes | 0.00675836 | 0.127818216 | -2.0422 | -1.03012123 |
| Tesc | 0.00199685 | 0.1078299 | -4.14128 | -2.050078159 |
| Tet1 | 0.00854669 | 0.133131133 | -2.26819 | -1.18153879 |
| Tgfb3 | 0.00822129 | 0.131828943 | -2.75812 | -1.463684454 |
| Tgif1 | 0.0016004 | 0.108284821 | 2.36149 | 1.239697426 |
| Tgm2 | 0.00106481 | 0.100624545 | -3.29684 | -1.721082711 |
| Thbs1 | 0.000032 | 0.09072 | 2.07884 | 1.055778724 |
| Thbs2 | 0.00375729 | 0.11368108 | 4.08534 | 2.030456151 |
| Thbs4 | 0.00421449 | 0.115106736 | 4.35911 | 2.12403361 |
| Thy1 | 0.00143508 | 0.107347013 | 3.11291 | 1.638263866 |
| Timp1 | 0.000766044 | 0.095251524 | 11.4413 | 3.51617908 |
| Timp4 | 0.00547856 | 0.122393362 | -10.8401 | -3.438308842 |
| Tinagl1 | 0.00185654 | 0.107195334 | -3.99191 | -1.997077177 |
| Tlr1 | 0.00988308 | 0.136942971 | 6.59819 | 2.722070322 |
| Tlr13 | 0.00285975 | 0.1112125 | 6.79306 | 2.764061597 |
| Tmem173 | 0.00540719 | 0.121952137 | 3.50664 | 1.810089328 |
| Tmem38b | 0.00085277 | 0.098276543 | -2.82015 | -1.495772176 |
| Tmem38b | 0.00288029 | 0.110795416 | -3.00419 | -1.58697791 |
| Tmod1 | 0.00182687 | 0.107007778 | -2.67023 | -1.416962376 |
| Tnc | 0.0016032 | 0.107958955 | 6.46676 | 2.693043069 |
| Tnfaip2 | 0.000574131 | 0.100472925 | 2.58943 | 1.372634559 |
| Tnfrsf1b | 0.00345546 | 0.112600334 | 2.17613 | 1.121764744 |
| Tnfrsf23 | 0.00847176 | 0.13298693 | 2.42082 | 1.275495812 |
| Tnfrsf26 | 0.00462704 | 0.116085473 | 2.51825 | 1.332421514 |
| Tnmd | 0.00643868 | 0.126497975 | 3.66181 | 1.872556936 |

| | | | | |
|---|---|---|---|---|
| Tnr | 0.00320554 | 0.1121939 | -5.36859 | -2.424540503 |
| Tns1 | 0.00321882 | 0.111967542 | -3.0487 | -1.608197093 |
| Tns1 | 0.00886225 | 0.133925793 | -3.93962 | -1.978054137 |
| Tns1 | 0.00888502 | 0.133984211 | -4.21127 | -2.074255733 |
| Tob2 | 0.00150413 | 0.107140918 | -2.08589 | -1.060662074 |
| Top2a | 0.00136596 | 0.106095797 | 9.06234 | 3.179883619 |
| Tpm1 | 0.00997799 | 0.137318455 | -2.03146 | -1.022516373 |
| Tpm2 | 0.0128218 | 0.147823518 | -3.36533 | -1.750751279 |
| Tpx2 | 0.0013001 | 0.105008077 | 3.32162 | 1.731887036 |
| Trim25 | 0.00863603 | 0.133132926 | 2.16534 | 1.114593574 |
| Trim36 | 0.00358627 | 0.112219376 | -2.18156 | -1.125361965 |
| Trim59 | 0.00172994 | 0.107788569 | 2.50664 | 1.325754813 |
| Trp53inp2 | 0.00175981 | 0.107755105 | -2.51589 | -1.331071232 |
| Trps1 | 0.00283251 | 0.111530081 | 2.78959 | 1.480053098 |
| Trps1 | 0.00350922 | 0.113052713 | 2.5614 | 1.356932568 |
| Tspan2 | 0.0112892 | 0.142497248 | -3.4372 | -1.781231229 |
| Tspan6 | 0.00895801 | 0.134655134 | 2.20301 | 1.139476044 |
| Ttc28 | 0.0000496 | 0.10044 | -2.25218 | -1.171319679 |
| Ttc28 | 0.000141684 | 0.11476404 | -2.45775 | -1.297338911 |
| Ttk | 0.0028824 | 0.110726341 | 2.6274 | 1.393635856 |
| Ttll7 | 0.00230762 | 0.108853622 | -2.65667 | -1.409619306 |
| Txlnb | 0.00133221 | 0.10520377 | -2.13927 | -1.097119123 |
| Tyrobp | 0.00800241 | 0.131137759 | 4.45247 | 2.154605891 |
| Uba2 | 0.00267563 | 0.110574505 | -3.00917 | -1.589367997 |
| Ugcg | 0.00356617 | 0.112710055 | 3.54863 | 1.827262159 |
| Unc93b1 | 0.00835434 | 0.132538074 | 2.64873 | 1.405300789 |
| Usp53 | 0.00097845 | 0.101237436 | -2.30412 | -1.204216431 |
| Vcl | 0.0102916 | 0.138278133 | -2.2012 | -1.138289141 |
| Vgll3 | 0.00195257 | 0.107070328 | -2.68105 | -1.422795011 |
| Vit | 0.000562735 | 0.101614887 | -5.40885 | -2.435323323 |
| Wdfy4 | 0.00325059 | 0.112246317 | 2.4981 | 1.32083123 |
| Wfdc1 | 0.00385472 | 0.114072351 | -3.0348 | -1.601601468 |
| Wfikkn2 | 0.0101986 | 0.138009695 | -2.60443 | -1.380964558 |
| Xirp1 | 0.00387816 | 0.113933509 | -2.81689 | -1.494100942 |
| Xk | 0.00112102 | 0.101213111 | -3.7271 | -1.898054155 |
| Zrsr1 | 0.00259464 | 0.111282971 | -2.24243 | -1.165065542 |
