## Supplementary material for "Survivin is a mechanosensitive cell cycle regulator in vascular smooth muscle cells": Table 2

**Table S2. Differentially expressed gene (DEG) lists of the *in vitro* study**

| Gene Symbol | p-value(stiff vs. soft) | q-values(stiff vs. soft) | Fold-Change(stiff vs. soft) | Log2FoldChange |
|---|---|---|---|---|
| 0610031J06Rik | 0.0000116 | 0.001949003 | -1.81205 | -0.857625775 |
| 1110018J18Rik | 0.00195994 | 0.032532069 | -1.73979 | -0.798910702 |
| 1700009N14Rik | 0.0020426 | 0.03336594 | 1.50567 | 0.590405607 |
| 1700029F09Rik | 0.0000841 | 0.00517808 | 1.59024 | 0.669244514 |
| 1810031K17Rik | 0.000115764 | 0.006043007 | -1.62577 | -0.701121192 |
| 2610002D18Rik | 0.00156593 | 0.028343136 | 1.89583 | 0.922829603 |
| 2610024B07Rik | 0.00000962 | 0.001898632 | 1.52103 | 0.605048608 |
| 2610036L11Rik | 0.000485955 | 0.013809461 | 2.15709 | 1.109086371 |
| 2810408A11Rik | 0.0000197 | 0.002551261 | -2.07371 | -1.052215665 |
| 2810417H13Rik | 0.000136022 | 0.006543362 | 3.38227 | 1.757991832 |
| 3110082D06Rik | 0.00000738 | 0.001687736 | 2.36453 | 1.241553446 |
| 4930422G04Rik | 0.0000394 | 0.003368875 | 1.77969 | 0.831625963 |
| 4930524J08Rik | 0.000449185 | 0.013113744 | 1.60052 | 0.678540705 |
| 5730469M10Rik | 0.000304549 | 0.010373025 | -1.53003 | -0.613564258 |
| 5730494M16Rik | 0.00116252 | 0.023507378 | 1.62432 | 0.699835879 |
| 5730590G19Rik | 0.00000144 | 0.001091937 | 2.38619 | 1.254708922 |
| 5730596B20Rik | 0.00144403 | 0.027036858 | -1.78931 | -0.839402454 |
| 6720463M24Rik | 0.00130917 | 0.025351971 | 1.58848 | 0.667646926 |
| 9030425E11Rik | 0.000097 | 0.005507498 | -1.62625 | -0.70155048 |
| 9930013L23Rik | 0.0025285 | 0.037967028 | 1.91328 | 0.936048021 |
| 9930111J21Rik | 0.00110206 | 0.022764057 | 1.8038 | 0.851039386 |
| A430107O13Rik | 0.0000692 | 0.004626445 | -2.38325 | -1.252934047 |
| A4galt | 0.0000821 | 0.005131695 | -1.59108 | -0.670008464 |
| A930001N09Rik | 0.000011 | 0.001962632 | -2.86124 | -1.516639767 |
| Abcc4 | 0.00312433 | 0.0434077 | -1.61806 | -0.694260562 |
| Abhd2 | 0.0000532 | 0.003851653 | -1.50839 | -0.593012796 |
| Abhd4 | 0.000654027 | 0.016373404 | -1.83683 | -0.877218628 |
| Acta2 | 0.000000155 | 0.000446633 | 1.56444 | 0.645646329 |
| Actg2 | 0.00000307 | 0.00138222 | 1.52937 | 0.61293748 |
| Adcyap1r1 | 0.00101399 | 0.021595064 | 1.61758 | 0.693837065 |
| Adk | 0.000380155 | 0.011893775 | 1.60448 | 0.682105806 |
| AI607873 | 0.000205989 | 0.008255317 | -3.18351 | -1.670616885 |
| Aif1l | 0.00179696 | 0.030894632 | 1.85115 | 0.888421803 |
| Ak3l1 | 0.00232844 | 0.036013955 | 1.66057 | 0.73167854 |
| Ak5 | 0.000390758 | 0.012107195 | 1.57823 | 0.658307469 |
| Akr1c13 | 0.000141817 | 0.00669911 | -1.5703 | -0.651035653 |
| Akr1c18 | 0.0000049 | 0.001551577 | -3.42637 | -1.776681254 |
| Alcam | 0.000197311 | 0.008064562 | -1.91878 | -0.940190663 |
| Aldh1a3 | 0.000534598 | 0.014601366 | -2.04455 | -1.031780932 |
| Amdhd2 | 0.000548588 | 0.014787243 | -1.625 | -0.700443505 |
| Angptl2 | 0.00000321 | 0.001284669 | -2.38846 | -1.256080092 |
| Ank | 0.0000483 | 0.003691683 | -2.23268 | -1.158777197 |
| Ankrd1 | 0.00000353 | 0.001320999 | 1.78908 | 0.8392179 |
| Ankrd26 | 0.00326978 | 0.044632265 | 1.67739 | 0.74621816 |
| Ankrd32 | 0.0000755 | 0.004845284 | 1.53704 | 0.62015471 |
| Ankrd37 | 0.000119372 | 0.00613138 | -1.72963 | -0.790459295 |

| | | | | |
|---|---|---|---|---|
| Anln | 0.00000724 | 0.001668965 | 3.2348 | 1.693676517 |
| Anpep | 0.0000114 | 0.001943734 | -2.29962 | -1.20139699 |
| Anxa3 | 0.000444379 | 0.013026227 | 1.60101 | 0.678982319 |
| Aox1 | 0.00384372 | 0.049400888 | -1.57484 | -0.655207855 |
| Apitd1 | 0.000735257 | 0.017684833 | 1.60972 | 0.686809763 |
| Appl2 | 0.0000271 | 0.002930156 | -1.82099 | -0.864725193 |
| Areg | 0.000454154 | 0.013205295 | -1.53931 | -0.622283188 |
| Arhgap11a | 0.000102575 | 0.005684036 | 2.26315 | 1.178332209 |
| Arhgap29 | 0.000975619 | 0.021010808 | 1.83652 | 0.876974607 |
| Arhgap6 | 0.0000497 | 0.003729441 | -2.31272 | -1.2095916 |
| Arrdc3 | 0.000100212 | 0.005596141 | -2.47696 | -1.308569463 |
| Asf1b | 0.0000541 | 0.003897229 | 2.00068 | 1.000490433 |
| Atad2 | 0.00010842 | 0.005806919 | 2.01016 | 1.007310338 |
| Atad5 | 0.00000103 | 0.000957402 | 2.20334 | 1.139692136 |
| Atp6ap2 | 0.0000997 | 0.00557836 | -1.5929 | -0.671650642 |
| Atp6v0b | 0.000821902 | 0.018946485 | -1.51495 | -0.599269724 |
| Atp6v0d1 | 0.000564769 | 0.014998911 | -1.51576 | -0.600041451 |
| Atp6v1b2 | 0.0000329 | 0.003098083 | -1.90612 | -0.930638787 |
| AU040829 | 0.00000179 | 0.001258021 | -1.57799 | -0.658084824 |
| Aurka | 0.000033 | 0.003097378 | 2.03307 | 1.023659889 |
| Aurkb | 0.0000732 | 0.004766685 | 1.94712 | 0.961341799 |
| AW549877 | 0.00329593 | 0.044882903 | -1.53837 | -0.621399598 |
| Bard1 | 0.000255287 | 0.009406771 | 2.48069 | 1.31074146 |
| BC031353 | 0.0000531 | 0.003868714 | -1.61248 | -0.689278314 |
| BC049807 | 0.000502752 | 0.014133462 | 1.72416 | 0.785893661 |
| BC064033 | 0.0000673 | 0.00456294 | -1.62602 | -0.701346376 |
| Bdh2 | 0.000816918 | 0.018892048 | -1.50849 | -0.593104197 |
| Bhlhe40 | 0.000536198 | 0.014617356 | -1.96083 | -0.971464794 |
| Birc5 | 0.000053 | 0.003876129 | 2.44717 | 1.291114326 |
| Blm | 0.000076 | 0.004839646 | 1.54242 | 0.625195664 |
| Blvrb | 0.00062916 | 0.015972903 | -1.7926 | -0.842050846 |
| Bmp2 | 0.000299781 | 0.010271331 | -3.44896 | -1.786163763 |
| Bpgm | 0.000107412 | 0.005806898 | -1.50814 | -0.592769088 |
| Brca1 | 0.00000346 | 0.001311841 | 2.10069 | 1.070863278 |
| Brca2 | 0.000111482 | 0.005894227 | 2.17634 | 1.12190396 |
| Brip1 | 0.0000153 | 0.002255087 | 2.82445 | 1.497969962 |
| Btc | 0.000125897 | 0.006309082 | -2.27549 | -1.186179824 |
| Bub1 | 0.0000432 | 0.003546462 | 3.88084 | 1.956368955 |
| Bub1b | 0.000288343 | 0.00999832 | 2.20146 | 1.138460631 |
| C330027C09Rik | 0.00184182 | 0.031366456 | 2.15646 | 1.108664956 |
| C3ar1 | 0.0000161 | 0.002325421 | 1.57288 | 0.653408607 |
| C4bp | 0.000442036 | 0.012997212 | 1.74522 | 0.803408912 |
| C79407 | 0.000112317 | 0.005927499 | 1.83881 | 0.878772417 |
| Calcoco1 | 0.0000852 | 0.005184874 | -1.52415 | -0.608003578 |
| Calcrl | 0.000603631 | 0.0157124 | -4.0292 | -2.010494733 |
| Camk1d | 0.000284329 | 0.009918814 | -1.57377 | -0.654222136 |
| Capg | 0.00000488 | 0.001571142 | -1.63521 | -0.709478869 |
| Car5b | 0.000614673 | 0.015842399 | -1.66111 | -0.732145436 |
| Casc5 | 0.0000264 | 0.002909048 | 3.08715 | 1.626275583 |

| | | | | |
|---|---|---|---|---|
| Cat | 0.0000371 | 0.003259258 | -1.77134 | -0.824839512 |
| Cby1 | 0.000267459 | 0.009694127 | -1.53094 | -0.614420973 |
| Ccdc15 | 0.000443664 | 0.013018511 | 1.51409 | 0.598450964 |
| Ccdc28b | 0.00176407 | 0.030621492 | -1.61953 | -0.695575406 |
| Ccdc85a | 0.000941948 | 0.020546731 | 1.5007 | 0.585635601 |
| Ccdc99 | 0.000197824 | 0.008051269 | 1.84031 | 0.879948809 |
| Cck | 0.00133068 | 0.025562363 | -1.50334 | -0.588165952 |
| Ccna2 | 0.0000298 | 0.0030396 | 2.9827 | 1.576618879 |
| Ccnb1 | 0.0000184 | 0.002448942 | 2.51784 | 1.332186608 |
| Ccne1 | 0.000179327 | 0.007666628 | 1.69741 | 0.763335082 |
| Ccne2 | 0.000630101 | 0.015954622 | 2.3717 | 1.245921533 |
| Ccng2 | 0.000350824 | 0.011396836 | -2.15246 | -1.105988625 |
| Ccnyl1 | 0.000014 | 0.002123211 | 2.16773 | 1.116185074 |
| Ccpg1 | 0.0000573 | 0.004076789 | -1.8713 | -0.904040483 |
| Cd3eap | 0.00000753 | 0.001681992 | 1.56839 | 0.649284348 |
| Cd68 | 0.000203337 | 0.00821761 | -1.5439 | -0.626580077 |
| Cd9 | 0.0000144 | 0.00217244 | -1.82891 | -0.870986091 |
| Cdc25c | 0.000183746 | 0.007752037 | 2.03809 | 1.027217761 |
| Cdc2a | 0.0000303 | 0.003021088 | 2.74943 | 1.459132556 |
| Cdc2l6 | 0.0000129 | 0.002036786 | -1.98326 | -0.987874751 |
| Cdc45l | 0.000317142 | 0.010663298 | 1.79038 | 0.840265825 |
| Cdc6 | 0.00000755 | 0.001673487 | 2.45746 | 1.297167934 |
| Cdc7 | 0.000120871 | 0.00615353 | 1.56165 | 0.64307115 |
| Cdca2 | 0.000150327 | 0.006908569 | 1.8893 | 0.917851804 |
| Cdca3 | 0.000551083 | 0.014840614 | 1.81728 | 0.861780722 |
| Cdca7 | 0.0000111 | 0.001956248 | 1.73488 | 0.794835876 |
| Cdca7l | 0.000638626 | 0.016127965 | 1.90865 | 0.932552572 |
| Cdca8 | 0.000119977 | 0.006140564 | 1.86732 | 0.900969181 |
| Cdk6 | 0.000279866 | 0.009858605 | 1.56291 | 0.644234703 |
| Cenpe | 0.0000954 | 0.00549241 | 2.53515 | 1.342071111 |
| Cenpf | 0.000134078 | 0.006482311 | 1.56408 | 0.645314306 |
| Cenph | 0.0000507 | 0.003774988 | 3.1343 | 1.648143274 |
| Cenpi | 0.000554396 | 0.014860391 | 1.77981 | 0.831723237 |
| Cenpk | 0.0000361 | 0.003240565 | 3.13896 | 1.650286645 |
| Cenpn | 0.000230211 | 0.008856515 | 2.07463 | 1.052854062 |
| Cenpp | 0.000293897 | 0.010129955 | 1.68725 | 0.754673754 |
| Cep55 | 0.000522212 | 0.014399559 | 2.92899 | 1.550403268 |
| Cep76 | 0.0000955 | 0.005476284 | 1.52723 | 0.610917348 |
| Chaf1a | 0.000243798 | 0.009135292 | 1.79537 | 0.844281193 |
| Chaf1b | 0.00020009 | 0.008120554 | 1.87997 | 0.91070964 |
| Chek1 | 0.0000912 | 0.005319692 | 2.46153 | 1.299555323 |
| Chic1 | 0.00221245 | 0.035066967 | -1.62011 | -0.696094199 |
| Ckap2 | 0.0000506 | 0.003777303 | 1.63566 | 0.709872891 |
| Ckap2l | 0.0000404 | 0.003403877 | 1.76976 | 0.823553727 |
| Cks2 | 0.0000754 | 0.004849667 | 1.51598 | 0.60025072 |
| Clspn | 0.00000558 | 0.001599878 | 2.52376 | 1.335574722 |
| Cmtm3 | 0.000106076 | 0.005799962 | -1.6173 | -0.69358609 |
| Cnn1 | 0.000195458 | 0.008045889 | 2.26903 | 1.182075684 |
| Col12a1 | 0.000148642 | 0.006863973 | 1.54761 | 0.630041956 |

| | | | | |
|---|---|---|---|---|
| Col6a1 | 0.0000628 | 0.004349957 | -1.62333 | -0.698953246 |
| Crabp1 | 0.000897503 | 0.019939513 | 2.20408 | 1.140176589 |
| Creb3l1 | 0.00199282 | 0.032775747 | 1.50333 | 0.588161734 |
| Creg1 | 0.000346946 | 0.011334749 | -1.58808 | -0.667281756 |
| Crim1 | 0.0000283 | 0.00299253 | 1.82099 | 0.864723 |
| Ctdspl2 | 0.000328093 | 0.01092948 | 1.54263 | 0.625392073 |
| Ctgf | 0.0000492 | 0.003701561 | 1.50675 | 0.591440065 |
| Ctns | 0.000165071 | 0.007363035 | -1.86363 | -0.898118681 |
| Ctps | 0.00042986 | 0.012915971 | 1.69802 | 0.763853452 |
| Ctsa | 0.00000665 | 0.001637776 | -1.74382 | -0.802247815 |
| Ctsb | 0.000122255 | 0.006191174 | -1.54132 | -0.624165394 |
| Ctsd | 0.00000583 | 0.001570014 | -1.53325 | -0.616595958 |
| Ctxn1 | 0.000217005 | 0.008542348 | -1.8855 | -0.914945243 |
| Cyp1a1 | 0.00222582 | 0.035124317 | -1.70787 | -0.772199788 |
| Cyp39a1 | 0.000830177 | 0.019000437 | -2.04813 | -1.034311006 |
| D0H4S114 | 0.000281624 | 0.009908419 | -1.62229 | -0.698035488 |
| D14Ertd436e | 0.0000632 | 0.004356718 | -1.62259 | -0.698299986 |
| D17H6S56E-5 | 0.00000482 | 0.00161498 | 1.83153 | 0.873049332 |
| Dab2 | 0.00000714 | 0.001659186 | -1.53169 | -0.615123506 |
| Dck | 0.00113941 | 0.023252195 | 1.51705 | 0.601268636 |
| Ddah2 | 0.00000446 | 0.001567255 | -1.71988 | -0.782307695 |
| Depdc1a | 0.000124675 | 0.006280612 | 1.88246 | 0.91261921 |
| Dhfr | 0.00363049 | 0.047746494 | 1.83232 | 0.873671481 |
| Dhrs3 | 0.0000667 | 0.004543642 | -2.39192 | -1.258166319 |
| Diap3 | 0.00000108 | 0.000928961 | 2.1158 | 1.08120326 |
| Dkc1 | 0.000187717 | 0.007850603 | 1.55296 | 0.63502067 |
| Dlc1 | 0.0000851 | 0.005195247 | 1.82631 | 0.868931671 |
| Dlgap5 | 0.0000148 | 0.002221156 | 3.01247 | 1.590946874 |
| Dna2 | 0.00000469 | 0.001608838 | 1.61126 | 0.688189312 |
| Dnajc9 | 0.000109481 | 0.005842028 | 1.52374 | 0.607616753 |
| Dnase1l1 | 0.000652523 | 0.016349957 | -1.58132 | -0.66112951 |
| Dnm1 | 0.0000413 | 0.003439478 | -1.72641 | -0.787776825 |
| Dnmt1 | 0.00000547 | 0.001592102 | 1.66443 | 0.735028197 |
| Dpp7 | 0.0000489 | 0.003698303 | -2.10858 | -1.076268153 |
| Dpt | 0.00204919 | 0.033397856 | -2.03553 | -1.025403143 |
| Dscc1 | 0.0000238 | 0.002770897 | 3.59021 | 1.844068233 |
| Dtl | 0.00000897 | 0.001872975 | 2.17307 | 1.119734648 |
| Dusp18 | 0.0000197 | 0.002551261 | -1.73304 | -0.79330927 |
| E2f7 | 0.000103626 | 0.005698441 | 1.68599 | 0.753595979 |
| E2f8 | 0.0000351 | 0.003170553 | 2.69523 | 1.430408392 |
| Ect2 | 0.000511826 | 0.014304817 | 2.35906 | 1.238212112 |
| Eif2b3 | 0.00163825 | 0.029211741 | 1.53693 | 0.620051458 |
| Emp2 | 0.00079351 | 0.018574322 | -1.53547 | -0.618683399 |
| Enpp1 | 0.000165104 | 0.007341777 | -1.79437 | -0.843481708 |
| Enpp3 | 0.000102165 | 0.005672224 | -3.64911 | -1.867541605 |
| Epha4 | 0.00224511 | 0.035274179 | 1.72776 | 0.788902829 |
| Epha7 | 0.000285922 | 0.00993829 | -1.81829 | -0.862577794 |
| Ephx1 | 0.00299514 | 0.04216168 | -1.53372 | -0.617034003 |
| Ercc6l | 0.00021184 | 0.008419544 | 2.84079 | 1.506292187 |

| | | | | |
|---|---|---|---|---|
| Esco2 | 0.00000312 | 0.001341833 | 1.81464 | 0.859683365 |
| Esm1 | 0.000138203 | 0.006615149 | -1.60549 | -0.683010904 |
| Exo1 | 0.0000167 | 0.002370495 | 2.9539 | 1.562620987 |
| Ezh2 | 0.00000941 | 0.001909501 | 1.56043 | 0.64194364 |
| F630043A04Rik | 0.00018816 | 0.007857725 | 2.29231 | 1.19680216 |
| F730047E07Rik | 0.000121502 | 0.006174744 | 2.04457 | 1.031797457 |
| Fabp3 | 0.00149954 | 0.027592111 | -1.83014 | -0.871952129 |
| Fabp4 | 0.001901 | 0.031921512 | -1.58807 | -0.667272592 |
| Fam100b | 0.0000372 | 0.003258109 | -1.68241 | -0.750532814 |
| Fam102a | 0.00114268 | 0.023285944 | -1.88818 | -0.916995008 |
| Fam102b | 0.00215232 | 0.034493382 | -1.5402 | -0.623120655 |
| Fam111a | 0.000167094 | 0.007373375 | 2.06838 | 1.04850126 |
| Fam150a | 0.000002 | 0.001340233 | 2.76651 | 1.468067139 |
| Fam33a | 0.000000237 | 0.000620832 | 1.51626 | 0.60051716 |
| Fam65b | 0.0000549 | 0.003935183 | 1.98982 | 0.99263793 |
| Fam98a | 0.0000456 | 0.003609791 | 1.52022 | 0.60428012 |
| Fancb | 0.00160802 | 0.028887217 | 2.04949 | 1.035264951 |
| Fancd2 | 0.00000154 | 0.001137823 | 1.6502 | 0.722640886 |
| Fancl | 0.00000316 | 0.001300791 | 1.84762 | 0.885668068 |
| Fbxl20 | 0.00013382 | 0.006480711 | -1.65321 | -0.72527198 |
| Fbxo5 | 0.000140286 | 0.006659541 | 2.69587 | 1.430750929 |
| Fen1 | 0.000175105 | 0.007564694 | 1.923 | 0.943358763 |
| Fez1 | 0.000512954 | 0.014294748 | 1.54385 | 0.626532588 |
| Fgfr1 | 0.00015628 | 0.007069401 | -1.63887 | -0.71269793 |
| Fgfr2 | 0.00076528 | 0.018149418 | 1.62693 | 0.702152179 |
| Fhl1 | 0.000364696 | 0.011663391 | 1.80541 | 0.852326503 |
| Fignl1 | 0.00000273 | 0.001404731 | 2.67166 | 1.41773642 |
| Fndc3a | 0.000000444 | 0.000609231 | -2.21804 | -1.149286973 |
| Fnip1 | 0.000218677 | 0.008573031 | -1.57711 | -0.6572837 |
| Foxm1 | 0.00081832 | 0.018909295 | 1.62809 | 0.703180453 |
| Foxq1 | 0.0000273 | 0.002935259 | -1.61065 | -0.687643843 |
| Fst | 0.00000254 | 0.001463802 | 1.65689 | 0.728477826 |
| Gadd45a | 3.25E-08 | 0.000312163 | -2.12619 | -1.088267007 |
| Gadd45g | 0.0000448 | 0.003595855 | 2.21468 | 1.147098258 |
| Gal3st4 | 0.000178017 | 0.007621931 | -1.51001 | -0.594558692 |
| Galm | 0.00211456 | 0.03413504 | -1.54265 | -0.625406726 |
| Galnt13 | 0.00114876 | 0.023327357 | 2.0994 | 1.06997707 |
| Gbe1 | 0.0000271 | 0.002930156 | -1.84195 | -0.881230982 |
| Gcnt1 | 0.000681108 | 0.016717314 | -2.1541 | -1.10708523 |
| Gda | 0.000824085 | 0.018936212 | -1.62317 | -0.698817418 |
| Gdpd1 | 0.000227058 | 0.008782116 | -1.5171 | -0.601316907 |
| Gem | 0.000402121 | 0.01232672 | -1.69724 | -0.763187805 |
| Gen1 | 0.00055393 | 0.014861725 | 2.31567 | 1.211429673 |
| Gins1 | 0.0000691 | 0.004630503 | 2.22682 | 1.154984946 |
| Gins2 | 0.0000217 | 0.002649515 | 1.79882 | 0.84705083 |
| Gla | 0.000000265 | 0.000636331 | -2.66422 | -1.413714674 |
| Gm10639 | 0.0000378 | 0.003280744 | -3.80043 | -1.926163318 |
| Gm12387 | 0.00000558 | 0.001599878 | 2.14207 | 1.099005626 |
| Gm14636 | 0.00214124 | 0.034373165 | -2.23945 | -1.163145142 |

| | | | | |
|---|---|---|---|---|
| Gm16494 | 0.000459615 | 0.013296994 | 1.75751 | 0.813532898 |
| Gm5465 | 0.0028015 | 0.040484063 | 2.15349 | 1.106676624 |
| Gm8074 | 0.000000685 | 0.000822428 | -1.71515 | -0.77833818 |
| Gm8681 | 0.000494737 | 0.013990036 | 1.60236 | 0.680198312 |
| Gm9948 | 0.00000417 | 0.001501982 | -2.17579 | -1.121536741 |
| Gmnn | 0.000829733 | 0.019005371 | 2.48906 | 1.315601008 |
| Gnpda1 | 0.000110808 | 0.00588017 | -1.64421 | -0.717394139 |
| Gpnmb | 0.0000814 | 0.005121269 | -2.40362 | -1.26520932 |
| Gpr137b | 0.000000795 | 0.000916317 | -2.30827 | -1.206811596 |
| Gpr137b-ps | 0.00000518 | 0.00158789 | -1.92549 | -0.945223746 |
| Gpr39 | 0.0000445 | 0.003601875 | 1.51281 | 0.597230805 |
| Grem1 | 0.000022 | 0.002663571 | -1.54244 | -0.625217565 |
| Grn | 0.00000852 | 0.001818547 | -1.96586 | -0.975155542 |
| Gsg2 | 0.0000387 | 0.00334877 | 2.22832 | 1.155956427 |
| Gsta1 | 0.0000156 | 0.002270273 | -4.88722 | -2.289016184 |
| Gsta2 | 0.0000459 | 0.003623585 | -3.70784 | -1.890577923 |
| Gsta3 | 0.0000102 | 0.001933638 | -3.21893 | -1.686582312 |
| Gsta4 | 0.000000951 | 0.000944933 | -6.41447 | -2.681325675 |
| Gstm1 | 0.000505958 | 0.014195891 | -1.94584 | -0.960395526 |
| Gstm3 | 0.000961679 | 0.020850851 | -1.6262 | -0.701503557 |
| Gstp2 | 0.0000227 | 0.00268074 | -1.52345 | -0.607339669 |
| Gtlf3a | 0.00281717 | 0.040608681 | 1.50213 | 0.587009675 |
| Gvin1 | 0.000608282 | 0.015762271 | 1.58931 | 0.668400554 |
| H2afv | 0.00000215 | 0.001376717 | -1.58112 | -0.660949293 |
| Hat1 | 0.00000485 | 0.001606353 | 1.81788 | 0.862256969 |
| Haus6 | 0.00156704 | 0.028345422 | 1.72809 | 0.789178356 |
| Haus7 | 0.0015055 | 0.02764881 | 1.62232 | 0.698058417 |
| Havcr2 | 0.00215318 | 0.034487983 | -1.83092 | -0.872567457 |
| Hbp1 | 0.000000413 | 0.000643275 | -2.10631 | -1.07472059 |
| Hells | 0.0000607 | 0.004266026 | 2.49758 | 1.32053089 |
| Herc3 | 0.000079 | 0.004981149 | -1.69273 | -0.759351065 |
| Hexb | 0.00002 | 0.002567038 | -1.76096 | -0.816364857 |
| Hist1h1b | 0.0000222 | 0.002670952 | 1.79053 | 0.840386691 |
| Hist1h2ab | 0.00222498 | 0.035130301 | 1.65543 | 0.727206008 |
| Hist1h2bc | 0.00310714 | 0.043294119 | 1.54604 | 0.628577646 |
| Hist1h2be | 0.00167414 | 0.029540933 | 1.68876 | 0.755964313 |
| Hist1h4f | 0.000950013 | 0.020691326 | 1.58033 | 0.660225849 |
| Hjurp | 0.000818978 | 0.018909336 | 1.66024 | 0.731391809 |
| Hmgb2 | 0.0000113 | 0.001955613 | 2.19403 | 1.133583253 |
| Hmgn2 | 0.000524486 | 0.014393394 | 1.56082 | 0.64230417 |
| Hmmr | 0.000329276 | 0.010956222 | 2.46437 | 1.301218878 |
| Hn1l | 0.000131391 | 0.006417003 | 1.67551 | 0.744600297 |
| Hoxa2 | 0.0000211 | 0.002598276 | -1.97944 | -0.985096301 |
| Hs6st2 | 0.0000675 | 0.004555064 | 1.88002 | 0.91074801 |
| Hsd17b11 | 0.000126714 | 0.006306155 | -2.31618 | -1.211748625 |
| Hsd3b7 | 0.000904207 | 0.019995951 | -1.53248 | -0.615868388 |
| Hspa4l | 0.000186526 | 0.007823503 | -1.51398 | -0.598349875 |
| Hspb2 | 0.000165316 | 0.007339878 | -1.50265 | -0.587513275 |
| Htr2a | 0.00199702 | 0.032807372 | 1.74859 | 0.806192054 |

| | | | | |
|---|---|---|---|---|
| Idh1 | 0.0000173 | 0.0024199 | -1.63466 | -0.708990615 |
| Idh3a | 0.00378858 | 0.04895423 | 1.53445 | 0.617721636 |
| Ifi47 | 0.00278784 | 0.040327113 | 1.69216 | 0.758865987 |
| Igf1 | 0.000422463 | 0.012760243 | 1.52057 | 0.604612233 |
| Il17rd | 0.000025 | 0.002836122 | -1.91261 | -0.93554202 |
| Il1rap | 0.000208017 | 0.008313467 | 1.57515 | 0.655489222 |
| Il1rl2 | 0.0014579 | 0.027172955 | -1.79066 | -0.840492228 |
| Incenp | 0.00000281 | 0.001372375 | 1.70274 | 0.767858159 |
| Ing4 | 0.000000945 | 0.000972506 | -1.53709 | -0.620199405 |
| Inpp5k | 0.000140472 | 0.006657402 | -1.72241 | -0.784430736 |
| Irgm1 | 0.00127642 | 0.024986442 | 1.54619 | 0.628717612 |
| Irs1 | 0.000563622 | 0.015009952 | 1.91236 | 0.935354135 |
| Itga11 | 0.00211884 | 0.034127655 | 1.51786 | 0.60203873 |
| Jam2 | 0.000571159 | 0.015112899 | -1.78895 | -0.839113363 |
| Jhdm1d | 0.0000152 | 0.002263504 | -2.55916 | -1.355667453 |
| Jmjd1c | 0.0000305 | 0.003014949 | -1.69707 | -0.763043345 |
| Kazald1 | 0.0000091 | 0.001872975 | -1.91839 | -0.939894495 |
| Kbtbd8 | 0.000235655 | 0.008970144 | 1.51015 | 0.594691857 |
| Kif11 | 0.0000101 | 0.00194021 | 3.13125 | 1.646738698 |
| Kif14 | 0.000409679 | 0.012478753 | 1.62723 | 0.702418182 |
| Kif15 | 0.000163598 | 0.007319994 | 2.35701 | 1.236957879 |
| Kif18a | 0.000245012 | 0.009145105 | 1.96542 | 0.974837642 |
| Kif20a | 0.0000298 | 0.0030396 | 1.61234 | 0.689156002 |
| Kif20b | 0.000367645 | 0.011705736 | 2.2973 | 1.199939268 |
| Kif22 | 0.00093358 | 0.02042605 | 1.52377 | 0.607645157 |
| Kif23 | 0.0000317 | 0.003049868 | 2.30358 | 1.203877702 |
| Kif2c | 0.000127438 | 0.006331252 | 1.53944 | 0.622405639 |
| Kif4 | 0.0000819 | 0.005130323 | 2.72556 | 1.44655268 |
| Klhl23 | 0.0000313 | 0.003062511 | 1.81864 | 0.86285999 |
| Klhl24 | 0.0000481 | 0.003696004 | -3.13059 | -1.646437315 |
| Kntc1 | 0.000018 | 0.00243507 | 2.59406 | 1.375211849 |
| Krt19 | 0.00279316 | 0.040383796 | 1.64821 | 0.720900069 |
| Lass6 | 0.00030148 | 0.010305037 | 1.79081 | 0.840612279 |
| Layn | 0.0000893 | 0.005294608 | -1.62538 | -0.700776446 |
| Lbr | 0.00000317 | 0.001286529 | 1.55286 | 0.634927768 |
| Lgals3 | 0.00000244 | 0.001434869 | -1.57254 | -0.653096424 |
| Lgmn | 0.00186371 | 0.031608478 | -1.57413 | -0.654555933 |
| Lig1 | 0.000025 | 0.002836122 | 1.61566 | 0.692123629 |
| Lin9 | 0.000619239 | 0.015903183 | 1.59827 | 0.676511147 |
| Lipa | 0.0000521 | 0.003839543 | -1.90135 | -0.927024385 |
| Lipg | 0.00137203 | 0.026044166 | -1.83454 | -0.875417761 |
| Lmnb1 | 6.47E-09 | 0.000186433 | 1.72644 | 0.787800196 |
| Lpin1 | 0.00114288 | 0.02327356 | -1.6473 | -0.720105615 |
| Ly96 | 0.000956062 | 0.020760306 | -1.67827 | -0.746974062 |
| Lyar | 0.0000208 | 0.002583414 | 1.54204 | 0.624840189 |
| Lyst | 0.00000333 | 0.001296675 | -1.55201 | -0.634139522 |
| Mad2l1 | 0.0000263 | 0.002920364 | 2.08181 | 1.057838405 |
| Mamdc2 | 0.0000181 | 0.002437157 | -2.92559 | -1.548725048 |
| Map1lc3b | 0.0000153 | 0.002255087 | -1.73244 | -0.792804307 |

| | | | | |
|---|---|---|---|---|
| Mastl | 0.0000247 | 0.002824327 | 2.5865 | 1.371001192 |
| Matn2 | 0.0000838 | 0.005176199 | -1.78673 | -0.837325901 |
| Mboat1 | 0.000258868 | 0.00949018 | 2.06819 | 1.048368729 |
| Mcm10 | 0.0000304 | 0.003020607 | 2.06659 | 1.047252194 |
| Mcm2 | 0.0000726 | 0.004776185 | 1.80799 | 0.854386698 |
| Mcm3 | 0.00000295 | 0.001371036 | 2.07425 | 1.052589786 |
| Mcm4 | 0.000000413 | 0.000643275 | 1.77627 | 0.828850894 |
| Mcm5 | 0.00000268 | 0.001404076 | 1.87109 | 0.903878954 |
| Mcm6 | 0.00000255 | 0.00144075 | 1.54959 | 0.631886549 |
| Mcm7 | 0.00000279 | 0.001386101 | 1.75957 | 0.815222909 |
| Mcm8 | 0.0000276 | 0.002956483 | 1.85457 | 0.891084723 |
| Mcpt8 | 0.00173872 | 0.030364374 | 2.04208 | 1.030039386 |
| Mef2c | 0.000207212 | 0.008292797 | -2.1338 | -1.093420291 |
| Megf10 | 0.000146958 | 0.006852095 | -1.72055 | -0.782871052 |
| Melk | 0.000179724 | 0.007660868 | 2.10614 | 1.074601339 |
| Mettl7a1 | 0.000325022 | 0.010839709 | -1.72367 | -0.785479751 |
| Mfsd11 | 0.000418774 | 0.012702077 | -1.55382 | -0.635815327 |
| Mgp | 0.00248057 | 0.037462067 | -1.64486 | -0.717963554 |
| Mgst1 | 0.00000675 | 0.001627626 | -2.30859 | -1.207014748 |
| Mgst2 | 0.0000977 | 0.005509248 | -1.96709 | -0.976066226 |
| Mki67 | 0.00000276 | 0.001395253 | 2.14844 | 1.103289487 |
| Mlf1 | 0.0000137 | 0.002088706 | 1.6657 | 0.736128588 |
| Mlf1ip | 0.0000107 | 0.00194524 | 2.50175 | 1.322937628 |
| Mmd | 0.000822497 | 0.018945045 | -1.5797 | -0.659651944 |
| Mmp10 | 0.00122398 | 0.024273217 | -1.75477 | -0.811280059 |
| Mmp11 | 0.000288477 | 0.009978949 | -1.69236 | -0.75903607 |
| Mmp13 | 0.00255096 | 0.038165064 | -1.5342 | -0.617487676 |
| Mmp19 | 0.0000119 | 0.001970681 | -1.81161 | -0.857275509 |
| Mns1 | 0.0000742 | 0.00481548 | 1.86823 | 0.901672078 |
| Morc4 | 0.00319627 | 0.044067234 | 1.69667 | 0.76270599 |
| Mr1 | 0.00369855 | 0.048245232 | -1.57808 | -0.658171336 |
| Mt1 | 0.0000126 | 0.002011463 | -1.91138 | -0.934617946 |
| Mtbp | 0.000282835 | 0.009890644 | 1.73337 | 0.793579641 |
| Mthfd1 | 0.0000359 | 0.003232683 | 1.68641 | 0.753955327 |
| Mthfd2 | 0.0000696 | 0.004642417 | 1.51958 | 0.603672629 |
| Mtmr11 | 0.000729097 | 0.017610168 | -1.65734 | -0.728870782 |
| Mybl1 | 0.00161028 | 0.028873813 | 1.76665 | 0.821016249 |
| Mybl2 | 0.00000711 | 0.001672446 | 1.86892 | 0.902204815 |
| Myl9 | 0.000252009 | 0.009333727 | 1.94938 | 0.963015348 |
| Myo19 | 0.00000646 | 0.001618651 | 1.69803 | 0.763861948 |
| Myo1e | 0.00000264 | 0.001421899 | -1.92658 | -0.946043457 |
| Myst4 | 0.0000869 | 0.005216716 | -1.6692 | -0.739155343 |
| Naga | 0.00195599 | 0.032485217 | -1.60304 | -0.680814477 |
| Nanos1 | 0.00071647 | 0.017378016 | 1.56279 | 0.644123929 |
| Nasp | 0.000132437 | 0.006435366 | 1.54435 | 0.626999751 |
| Nav1 | 0.000540125 | 0.014668899 | -1.57175 | -0.652368357 |
| Ncam1 | 0.00000745 | 0.001690329 | 2.02267 | 1.016260962 |
| Ncapd3 | 0.000195532 | 0.008037453 | 1.59863 | 0.676836068 |
| Ncapg | 0.0000297 | 0.003056448 | 3.36097 | 1.748877665 |

| | | | | |
|---|---|---|---|---|
| Ncapg2 | 0.00000891 | 0.001874027 | 2.59292 | 1.374577695 |
| Ncaph | 0.000672476 | 0.016632958 | 2.04539 | 1.032375952 |
| Ndc80 | 0.000389197 | 0.012097855 | 2.56541 | 1.359189414 |
| Ndrg2 | 0.0000831 | 0.005160617 | -2.22904 | -1.156424508 |
| Nek7 | 0.000156208 | 0.007077254 | 1.53147 | 0.614917106 |
| Nfatc1 | 0.000479287 | 0.013687468 | -1.76526 | -0.819877606 |
| Nfatc4 | 0.0000268 | 0.002914121 | -1.75462 | -0.811161079 |
| Nfib | 0.00257968 | 0.03837557 | 1.52468 | 0.608506481 |
| Ngef | 0.00149235 | 0.027512518 | -1.56768 | -0.64862948 |
| Nipal1 | 0.0012515 | 0.024632495 | 1.5773 | 0.657457085 |
| Nme5 | 0.00106604 | 0.022259379 | -1.84233 | -0.881533953 |
| Nol10 | 0.0000665 | 0.004540752 | 1.5027 | 0.587557017 |
| Nov | 0.00207586 | 0.033699102 | -1.57882 | -0.658849949 |
| Npc1 | 0.000276252 | 0.009827409 | -1.57157 | -0.652202835 |
| Npnt | 0.0000341 | 0.003134263 | 2.38251 | 1.25248227 |
| Nqo1 | 0.000037 | 0.003270414 | -2.81494 | -1.493105629 |
| Nr1d1 | 0.0000238 | 0.002770897 | -1.62608 | -0.70139564 |
| Nr2f1 | 0.0000174 | 0.00242213 | -1.7539 | -0.810563796 |
| Nsl1 | 0.0000454 | 0.00360883 | 1.67247 | 0.741980333 |
| Nt5dc2 | 0.000261925 | 0.009553631 | 1.8319 | 0.873340752 |
| Nuf2 | 0.000105295 | 0.005768204 | 3.89587 | 1.961945537 |
| Nup107 | 0.000135799 | 0.006543559 | 1.64432 | 0.717491088 |
| Nup160 | 0.000061 | 0.004266299 | 1.5139 | 0.598269912 |
| Nup43 | 0.0000705 | 0.004691588 | 1.87934 | 0.910226095 |
| Nup85 | 0.00000264 | 0.001421899 | 1.60658 | 0.683992822 |
| Nusap1 | 0.0000527 | 0.003863996 | 2.29772 | 1.200203002 |
| Ociad2 | 0.000006 | 0.001593456 | -1.75838 | -0.814247607 |
| Ogfrl1 | 0.000146304 | 0.006854878 | 1.62552 | 0.700901307 |
| Olfml3 | 0.00248774 | 0.03755067 | -1.6334 | -0.707880276 |
| Olr1 | 0.000646569 | 0.016257317 | -1.9688 | -0.97731545 |
| Orc1l | 0.00000301 | 0.001376717 | 2.01654 | 1.011882023 |
| Orc6l | 0.0000184 | 0.002448942 | 1.5872 | 0.666483931 |
| P2rx4 | 0.00101308 | 0.021591642 | -1.70031 | -0.765795465 |
| Pamr1 | 0.0000265 | 0.002897903 | -3.63138 | -1.860520027 |
| Pard6b | 0.000125527 | 0.006312497 | 1.50796 | 0.59259816 |
| Pask | 0.0000161 | 0.002325421 | 1.74587 | 0.803946138 |
| Pbk | 0.00000172 | 0.001239045 | 3.60992 | 1.851966866 |
| Pcdh18 | 0.0000253 | 0.002847732 | 1.69288 | 0.759479711 |
| Pcdhb16 | 0.0000205 | 0.002579509 | -1.79342 | -0.84271047 |
| Pcdhb20 | 0.000191348 | 0.007933371 | -1.58056 | -0.660431582 |
| Pcmtd1 | 0.000349416 | 0.011376748 | -1.61127 | -0.688194659 |
| Pcmtd2 | 0.000378421 | 0.011891168 | -1.99877 | -0.99910869 |
| Pcsk6 | 0.0000919 | 0.005338908 | 1.52919 | 0.612767671 |
| Pdlim5 | 6.85E-08 | 0.000328971 | 1.53195 | 0.615369211 |
| Peg10 | 0.000000854 | 0.000911408 | 1.75009 | 0.807429116 |
| Penk | 0.00213131 | 0.034251923 | -1.92436 | -0.944376738 |
| Pip4k2c | 0.0010545 | 0.022114569 | -1.50151 | -0.586418915 |
| Plac8 | 0.000174732 | 0.007571282 | 1.54584 | 0.628391003 |
| Plau | 0.00278229 | 0.040267045 | 1.55059 | 0.632817266 |

| | | | | |
|---|---|---|---|---|
| Plcb4 | 0.0000575 | 0.004080942 | 1.50649 | 0.591191096 |
| Plekhm1 | 0.000499446 | 0.01409553 | -1.54664 | -0.629132682 |
| Plin2 | 0.00000457 | 0.001586561 | -2.01468 | -1.010552863 |
| Plk1 | 0.00128697 | 0.025124689 | 1.68565 | 0.753305013 |
| Plk4 | 0.000521478 | 0.014393093 | 1.92424 | 0.94428875 |
| Plxna4 | 0.0000913 | 0.005314767 | -1.64089 | -0.714477044 |
| Pms1 | 0.000000614 | 0.0008042 | 1.55049 | 0.632724221 |
| Pms2 | 0.000349616 | 0.011370412 | 1.65978 | 0.730992028 |
| Pola1 | 0.0000663 | 0.004537849 | 1.7634 | 0.818359765 |
| Pola2 | 0.00103988 | 0.022000104 | 1.50366 | 0.588478389 |
| Pole | 0.000000147 | 0.000470645 | 2.35746 | 1.237233292 |
| Pole2 | 0.000152899 | 0.006960165 | 1.74149 | 0.800322189 |
| Polh | 0.0000443 | 0.003595787 | 1.69469 | 0.761021394 |
| Polq | 0.00012641 | 0.006312832 | 1.51149 | 0.595971434 |
| Pop1 | 0.003636 | 0.047731818 | 1.51328 | 0.597678952 |
| Ppil1 | 0.000805023 | 0.018782784 | 1.56204 | 0.643431398 |
| Ppil5 | 0.0000283 | 0.00299253 | 2.49153 | 1.317031945 |
| Prc1 | 0.0000761 | 0.004830003 | 2.40523 | 1.266174858 |
| Prdm1 | 0.00175068 | 0.030499301 | -1.56749 | -0.648453079 |
| Prex2 | 0.000502728 | 0.014146589 | 1.73698 | 0.796581143 |
| Prim1 | 0.0000131 | 0.002051503 | 2.77068 | 1.470240096 |
| Prim2 | 0.000117074 | 0.00605653 | 1.95938 | 0.970397219 |
| Prkd1 | 0.00169578 | 0.029776905 | 1.63151 | 0.70620783 |
| Prl2c3 | 0.0000346 | 0.00315506 | -2.04853 | -1.034585831 |
| Prl2c5 | 0.000006 | 0.001593456 | -2.33715 | -1.224748823 |
| Prps1 | 0.000386348 | 0.012035262 | 1.52817 | 0.611805044 |
| Prr11 | 0.000333945 | 0.011073217 | 2.67953 | 1.421979969 |
| Prrx2 | 0.0030664 | 0.042871575 | -1.50026 | -0.585214994 |
| Prss23 | 0.000204473 | 0.008228896 | 1.54811 | 0.630507985 |
| Psap | 0.000747211 | 0.017853138 | -1.52777 | -0.611426898 |
| Psat1 | 0.000106743 | 0.005792466 | 1.6372 | 0.711230572 |
| Psmc3ip | 0.0000823 | 0.005133062 | 1.83612 | 0.876660349 |
| Ptgr1 | 0.00002 | 0.002567038 | -1.88352 | -0.913433605 |
| Ptgs1 | 0.0001032 | 0.005696759 | -1.66338 | -0.734114282 |
| Ptprk | 0.000282823 | 0.009902241 | 1.56498 | 0.64614422 |
| Ptprm | 0.00000108 | 0.000928961 | -1.5088 | -0.59340238 |
| Pus7l | 0.00039183 | 0.012114358 | 1.64458 | 0.717719189 |
| Rab31 | 0.000277661 | 0.009822961 | -1.68665 | -0.754161198 |
| Rab5b | 0.000284415 | 0.009909816 | -1.52888 | -0.612476433 |
| Racgap1 | 0.0000306 | 0.003009348 | 2.04056 | 1.028965132 |
| Rad51 | 0.00000943 | 0.001900178 | 3.01436 | 1.591851726 |
| Rad51ap1 | 0.000000325 | 0.000624325 | 1.96455 | 0.974198886 |
| Rad54b | 0.00018157 | 0.00770536 | 1.62298 | 0.698645222 |
| Rad54l | 0.00000638 | 0.0016269 | 2.01683 | 1.012089483 |
| Rbl1 | 0.0000104 | 0.001927177 | 2.14749 | 1.102651413 |
| Rbp1 | 0.00000949 | 0.001898989 | -1.93512 | -0.952422526 |
| Rcan2 | 0.000072 | 0.00475844 | -1.77591 | -0.828557441 |
| Renbp | 7.01E-09 | 0.000100997 | -2.47783 | -1.309076988 |
| Rfc2 | 0.000511961 | 0.014294725 | 1.75944 | 0.815116316 |

| | | | | |
|---|---|---|---|---|
| Rfc3 | 0.0000299 | 0.003012477 | 1.72247 | 0.784478856 |
| Rfc4 | 0.000316727 | 0.010661786 | 2.03326 | 1.023794709 |
| Rftn2 | 0.000231276 | 0.008873792 | -1.59051 | -0.669489786 |
| Rgs16 | 0.000376866 | 0.01186819 | 1.52408 | 0.607938633 |
| Rin2 | 0.000224485 | 0.008717703 | -1.5128 | -0.597218894 |
| Rnd1 | 0.00277924 | 0.040263349 | 2.03319 | 1.02374504 |
| Rnf115 | 0.00000571 | 0.001582054 | -1.68273 | -0.750799832 |
| Rnu2 | 0.00373447 | 0.048516119 | -1.53039 | -0.613899818 |
| RP23-38E20.1 | 0.00148985 | 0.027554575 | 1.64531 | 0.718359434 |
| Rpa2 | 0.000185379 | 0.007798096 | 1.63879 | 0.712630995 |
| Rpp30 | 0.00286496 | 0.040989981 | 1.5137 | 0.598079306 |
| Rras2 | 0.0000751 | 0.004857478 | 1.55024 | 0.632491583 |
| Rrm1 | 0.0000894 | 0.005289653 | 1.9207 | 0.941632198 |
| Rrm2 | 0.0000316 | 0.003065838 | 2.22695 | 1.155069167 |
| Rrp1b | 0.000284496 | 0.009900667 | 1.56867 | 0.649541885 |
| S1pr1 | 0.0000401 | 0.003398475 | 1.99689 | 0.997754863 |
| Samhd1 | 0.0000323 | 0.003076775 | 1.82073 | 0.864516998 |
| Sash1 | 0.000128152 | 0.006323116 | -1.5799 | -0.659831998 |
| Sat1 | 0.0000336 | 0.003133282 | -1.58084 | -0.660686993 |
| Scel | 0.000597205 | 0.015573269 | 1.98163 | 0.986687615 |
| Scyl3 | 0.000548467 | 0.014797825 | 1.57965 | 0.659604939 |
| Sdpr | 0.000188636 | 0.007854836 | 1.97123 | 0.979096118 |
| Sema3a | 0.000742087 | 0.017789715 | 1.50703 | 0.591708137 |
| Sema6d | 0.0000517 | 0.003819835 | -2.18919 | -1.130397027 |
| Sepp1 | 0.000165359 | 0.007330492 | -1.81582 | -0.860616951 |
| Serpinb1a | 0.00253291 | 0.03799365 | -1.75769 | -0.813682009 |
| Serpinb8 | 0.00211191 | 0.034111371 | -1.59845 | -0.676669134 |
| Serpine2 | 0.0000114 | 0.001943734 | -2.99056 | -1.580417963 |
| Sgol1 | 0.0000852 | 0.005184874 | 2.03876 | 1.027691953 |
| Sgol2 | 0.00134097 | 0.025640379 | 2.02486 | 1.017822163 |
| Sgpl1 | 0.000005 | 0.001566033 | -1.57667 | -0.656878755 |
| Shcbp1 | 0.00012814 | 0.006333369 | 2.98969 | 1.5799959 |
| Skp2 | 0.000171548 | 0.00748963 | 1.55025 | 0.632500889 |
| Slc14a1 | 0.0000365 | 0.003261233 | -2.75736 | -1.463290574 |
| Slc16a1 | 0.000364384 | 0.011666361 | 1.60355 | 0.681269339 |
| Slc17a5 | 0.000236243 | 0.008980662 | -1.89748 | -0.924086808 |
| Slc1a6 | 0.000160742 | 0.007237157 | 1.88266 | 0.912772479 |
| Slc40a1 | 0.000021 | 0.00259706 | -1.90623 | -0.930718538 |
| Slc44a1 | 0.000603929 | 0.015691807 | -1.53305 | -0.616405737 |
| Slc4a4 | 0.000212814 | 0.008434987 | 2.05799 | 1.041235972 |
| Slfn9 | 0.0000317 | 0.003049868 | 2.02212 | 1.015868615 |
| Smc2 | 0.0000288 | 0.003023213 | 2.1721 | 1.119090524 |
| Smc4 | 0.00276999 | 0.040210207 | 1.64356 | 0.716824125 |
| Snx24 | 0.000161237 | 0.007236829 | 1.54243 | 0.625205017 |
| Sorbs1 | 0.0000423 | 0.003502513 | 1.58301 | 0.662670369 |
| Sox11 | 0.00348274 | 0.046482239 | 1.5383 | 0.621336886 |
| Sox9 | 0.0000729 | 0.004779553 | -2.28312 | -1.191007107 |
| Spag5 | 0.00331448 | 0.04507161 | 1.61952 | 0.695566285 |
| Spc24 | 0.000166504 | 0.007369912 | 2.40067 | 1.263437102 |

| | | | | |
|---|---|---|---|---|
| Spc25 | 0.0000896 | 0.005290623 | 2.41949 | 1.274702976 |
| Speer3 | 0.00237898 | 0.03646293 | 1.54565 | 0.62821367 |
| Speer4a | 0.00378537 | 0.048956659 | -2.11243 | -1.078901912 |
| Spp1 | 0.000748977 | 0.017880507 | -2.66693 | -1.415179852 |
| Spred1 | 0.00149136 | 0.027529493 | -1.55257 | -0.634661322 |
| Spred2 | 0.000250562 | 0.009292077 | -1.55547 | -0.637351697 |
| Sprr2h | 0.0000963 | 0.005494821 | 2.56757 | 1.360403609 |
| Sqle | 0.00115069 | 0.023317252 | 1.61897 | 0.695076252 |
| Sqstm1 | 0.0000214 | 0.002624004 | -1.84817 | -0.886099511 |
| Srgap1 | 0.000458133 | 0.01326744 | -1.57312 | -0.653625127 |
| Ssbp2 | 0.0000219 | 0.002662652 | -1.65417 | -0.726109386 |
| Ssx2ip | 0.0000226 | 0.002679914 | 1.549 | 0.631337144 |
| Star | 0.000866702 | 0.019495721 | -1.64559 | -0.718606788 |
| Stard4 | 0.00149876 | 0.02759538 | 1.60215 | 0.680009225 |
| Stil | 0.00132953 | 0.02555731 | 1.83921 | 0.879086215 |
| Stom | 0.000471299 | 0.013499484 | -1.60119 | -0.67914336 |
| Suv39h2 | 0.0000396 | 0.003366 | 1.76996 | 0.823716757 |
| Tacc3 | 0.00000559 | 0.001579175 | 2.17805 | 1.123037073 |
| Tagln | 0.00000815 | 0.001765731 | 2.04573 | 1.032615748 |
| Tarsl2 | 0.0000316 | 0.003065838 | 1.77126 | 0.824775998 |
| Tcf19 | 0.000493518 | 0.013969274 | 1.85094 | 0.88825813 |
| Tcn2 | 0.0000654 | 0.004497616 | -1.78744 | -0.837895688 |
| Tcp11l2 | 0.000222108 | 0.008637034 | -2.41292 | -1.270782248 |
| Tdrkh | 0.00208323 | 0.033799703 | 1.67603 | 0.745047973 |
| Tead1 | 0.0000126 | 0.002011463 | 1.55285 | 0.634918477 |
| Tfrc | 0.0000391 | 0.003363184 | 1.67432 | 0.743575286 |
| Tgfb1 | 0.0000116 | 0.001949003 | -1.82188 | -0.865432066 |
| Tgfb2 | 0.000864043 | 0.019466301 | 1.65159 | 0.723855588 |
| Tgfbi | 0.0000343 | 0.003137633 | -4.37208 | -2.128320339 |
| Tgm2 | 0.000700216 | 0.017055557 | 1.58526 | 0.664719478 |
| Tipin | 0.00000397 | 0.001448045 | 1.66466 | 0.735227543 |
| Tlr4 | 0.000407804 | 0.012447958 | -1.6575 | -0.729009469 |
| Tmem140 | 0.0000169 | 0.002387125 | -2.53844 | -1.343944857 |
| Tmem48 | 0.000454359 | 0.013197938 | 1.62357 | 0.699169588 |
| Tmem50a | 0.0000291 | 0.003038103 | -1.56811 | -0.649025328 |
| Tmem59 | 0.000028 | 0.002988222 | -1.54312 | -0.625854134 |
| Tmem77 | 0.000177508 | 0.007611448 | -1.57021 | -0.650960894 |
| Tmem97 | 0.000446284 | 0.013068774 | 1.7617 | 0.816968269 |
| Tnfaip2 | 0.0000476 | 0.003682132 | -1.88212 | -0.912355221 |
| Tnfrsf22 | 0.0000117 | 0.00194876 | -1.63846 | -0.712338589 |
| Tnfrsf23 | 0.0000627 | 0.004358747 | -1.72267 | -0.784649425 |
| Tnfsf15 | 0.000557751 | 0.014894898 | 2.07951 | 1.056243623 |
| Top2a | 0.0000053 | 0.001607574 | 3.07097 | 1.618694419 |
| Topbp1 | 0.000118175 | 0.006080737 | 1.55231 | 0.634416696 |
| Tpp1 | 0.0000847 | 0.005192831 | -1.76781 | -0.821959783 |
| Tpx2 | 0.0000337 | 0.003132469 | 1.87117 | 0.903940637 |
| Trip13 | 0.000131371 | 0.006426919 | 2.22303 | 1.152527418 |
| Trp53inp1 | 0.000049 | 0.003696165 | -1.76352 | -0.818454688 |
| Tshz1 | 0.000843099 | 0.019219856 | -1.71183 | -0.775537355 |

| | | | | |
|---|---|---|---|---|
| Tspan12 | 0.00006 | 0.004227139 | -3.45142 | -1.787194122 |
| Tspan2 | 0.000147345 | 0.006847978 | 1.55478 | 0.636710455 |
| Ttc33 | 0.000341942 | 0.011234959 | -1.50731 | -0.59197297 |
| Ttc9 | 0.00166968 | 0.029498362 | -1.52683 | -0.610536714 |
| Ttk | 0.00000574 | 0.001560359 | 3.67046 | 1.87596088 |
| Tyms | 0.0000393 | 0.003370326 | 1.77174 | 0.825166906 |
| U90926 | 0.0000123 | 0.002002398 | 3.21827 | 1.686285367 |
| Uap1l1 | 0.0000446 | 0.003599857 | -1.79669 | -0.845341619 |
| Ube2c | 0.000969806 | 0.020916886 | 1.59405 | 0.672696883 |
| Ube2t | 0.00000309 | 0.001369821 | 1.68983 | 0.756878116 |
| Uhrf1 | 0.000101557 | 0.00566028 | 2.46476 | 1.301447175 |
| Umps | 0.00257496 | 0.038364774 | 1.56348 | 0.644760764 |
| Usp1 | 0.0000136 | 0.002101255 | 1.63289 | 0.707427607 |
| Usp37 | 0.000174877 | 0.007566187 | 1.58071 | 0.660572712 |
| Vgll3 | 0.000363179 | 0.011653678 | 1.56242 | 0.643782322 |
| Vrk1 | 0.00000227 | 0.001421958 | 1.73962 | 0.7987722 |
| Vwa5a | 0.0000177 | 0.002428693 | -1.85071 | -0.88807663 |
| Wbp2 | 0.0000302 | 0.003021573 | -1.5926 | -0.671381794 |
| Wdhd1 | 0.0000136 | 0.002101255 | 2.41062 | 1.269404248 |
| Wdr19 | 0.00122329 | 0.024276241 | -1.5563 | -0.638119373 |
| Wdr45 | 0.000892962 | 0.0199 | -1.56943 | -0.650236168 |
| Wdr76 | 0.0000591 | 0.004173938 | 2.08197 | 1.05794928 |
| Wee1 | 0.000759383 | 0.018069051 | 1.61431 | 0.69091765 |
| Wfdc3 | 0.000000337 | 0.000606916 | -1.76098 | -0.8163801 |
| Xdh | 0.00020043 | 0.008122912 | -1.98647 | -0.990211415 |
| Ypel3 | 0.0000558 | 0.003979894 | -1.59734 | -0.675673256 |
| Zfp260 | 0.00062351 | 0.015955986 | -1.56727 | -0.64825635 |
| Zwilch | 0.00186284 | 0.031612329 | 2.13557 | 1.094621188 |
